# Epigenetic modulators link mitochondrial redox homeostasis to cardiac function

**DOI:** 10.1101/2022.03.26.485908

**Authors:** Zaher Elbeck, Mohammad Bakhtiar Hossain, Humam Siga, Nikolay Oskolkov, Fredrik Karlsson, Julia Lindgren, Anna Walentinsson, Cristobal Dos Remedios, Dominique Koppenhöfer, Rebecca Jarvis, Roland Bürli, Tanguy Jamier, Elske Franssen, Mike Firth, Andrea Degasperi, Claus Bendtsen, Jan Dudek, Michael Kohlhaas, Alexander G. Nickel, Lars H. Lund, Christoph Maack, Ákos Végvári, Christer Betsholtz

## Abstract

Excessive production of reactive oxygen species (ROS) is characteristic of numerous diseases, but most studies in this area have not considered the impact of endogenous antioxidative defenses. Here, utilizing multi-omics, we demonstrate that in cardiomyocytes mitochondrial isocitrate dehydrogenase (IDH2) constitutes a major antioxidant defense. In both male and female mice and humans the paradoxical reduction in expression of IDH2 associated with heart failure is compensated for by an increase in the enzyme’s activity. We describe extensive mutual regulation of the antioxidant activities of IDH2 and NRF2 by a network involving 2-oxoglutarate and L2-hydroxyglutarate and mediated in part through unconventional hydroxymethylation of cytosine residues present in introns. Conditional targeting of ROS in a murine model of heart failure improves cardiac function. Together, these insights may explain why previous attempts to treat heart failure with antioxidants have been unsuccessful and open new approaches to personalizing and, thereby, improving such treatment.

**Graphical abstract:** 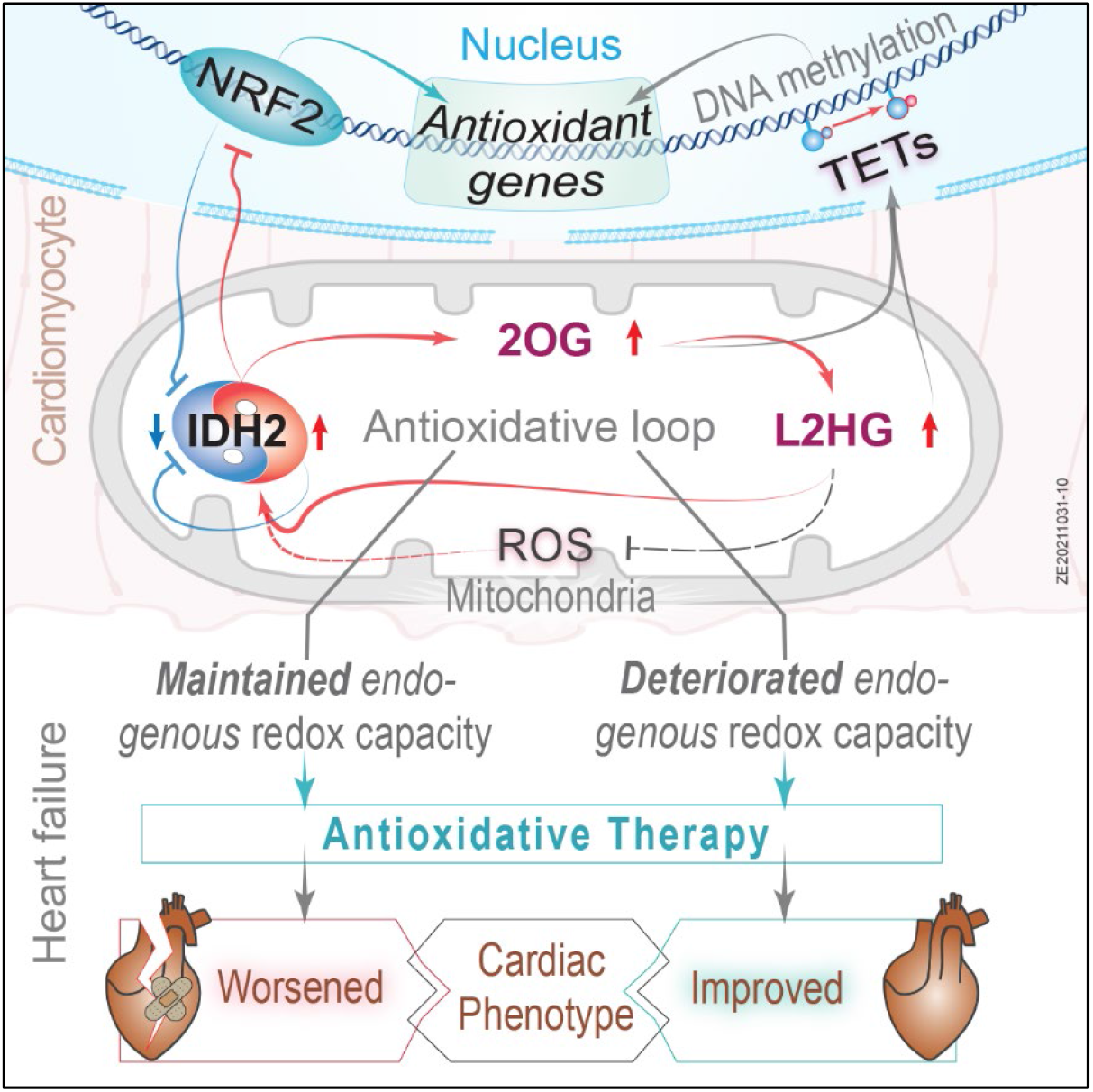

**Highlights:**

- Paradoxical downregulation of mitochondrial isocitrate dehydrogenase (IDH2) in response to oxidative stress leads to the discovery of a robust antioxidative defense in the heart.
- An antioxidative loop involving IDH2 coordinates other antioxidative defenses, such as NRF2.
- This loop produces epigenetic modifications that link oxidative stress to mitochondrial function.
- The conclusion that enhancing antioxidative capacity improves cardiac function only when the endogenous capacity is insufficient opens new approaches to individualized treatment of patients with heart failure.

## Introduction

The incidence of heart failure continues to increase worldwide and despite improvements in medical and device-based treatments, morbidity and mortality remain unacceptably high. In this context, metabolic dysfunction and oxidative stress play crucial roles (Bertero and Maack, 2018b; Munzel et al., 2017), with mitochondria being the major source of reactive oxygen species (ROS) in cardiac tissue. Indeed, formation of ROS through leakage of electrons from complex I of the respiratory chain is elevated in failing hearts (Ide et al., 2000; Ide et al., 1999).

More recently, disturbance of the coupling between excitation and contraction in failing cardiac myocytes has been shown to attenuate activation of the Krebs cycle by Ca^2+^, causing oxidation of NADH, FADH_2_ and NADPH and thereby disrupting enhancement of energy production in response to greater demand (Bertero and Maack, 2018a). While NADH and FADH_2_ donate electrons to the mitochondrial respiratory chain for production of ATP, NADPH is a necessary cofactor for antioxidative processes that eliminate hydrogen peroxide (H_2_O_2_), i.e., the enzyme glutathione peroxidase and the peroxiredoxin/thioredoxin system (Nickel et al., 2014). Nonetheless, clinical trials designed to treat heart failure by ameliorating oxidative stress have, to date, been unsuccessful (van der Pol et al., 2019; Zhou and Tian, 2018) and, in some patients with chronic renal disease, even been reported to promote the development of heart failure (de Zeeuw et al., 2013).

One potential explanation for this failure is that signaling by physiological levels of ROS has protective effects on the heart and, consequently, excessive quenching of these species may be undesirable (Santos et al., 2011; Song et al., 2014). Accordingly, in the attempt to develop more effective treatments of heart failure, a more detailed molecular understanding of cardiac redox regulation is necessary. Furthermore, since heart failure is associated with a variety of etiologies and phenotypes (Triposkiadis et al., 2019), an individualized approach to interventions that alter mitochondrial metabolism and redox status in cardiac tissue may be required.

Alterations in mitochondrial biogenesis and function in response to various stimuli and forms of stress are tightly regulated through control of the expression of key nuclear genes. In addition to producing ATP, it is becoming increasingly clear that mitochondria generate metabolites that participate in cellular signal transduction, as well as serve as cofactors for various biochemical processes, including several reactions involved in gene expression and its regulation (Martínez-Reyes and Chandel, 2020). For instance, 2-oxoglutarate (2OG, also known as α-ketoglutarate, α-KG) produced via the Krebs cycle is an essential cofactor for certain dioxygenases (OGDD) that catalyze hydroxylation of nucleic acids, chromatin, proteins, lipids, and metabolites (Martínez-Reyes and Chandel, 2020). A major route for synthesis of 2OG involves oxidative decarboxylation of isocitrate catalyzed by mitochondrial isocitrate dehydrogenase (IDH2).

Importantly, IDH2 plays an essential role in antioxidant systems, is present at high levels in cardiac tissue (Calvo et al., 2016), and is mainly responsible for direct regeneration of mitochondrial NADPH in cardiac myocytes (Nickel et al., 2015). In addition, IDH2 contributes indirectly to NADPH production by providing the substrate for 2-oxoglutarate dehydrogenase (OGDH), which generates NADH that is utilized preferentially by nicotinamide nucleotide transhydrogenase (NNT) to regenerate NADPH (Wagner et al., 2020). The activity of IDH2 is altered through changes in NAD^+^ levels or oxidative stimuli, which are modulated by SIRT3 and SIRT5 (Yu et al., 2012; Zhou et al., 2016), as well as stimulated when the Krebs cycle is activated by ADP and/or Ca^2+^ (Kohlhaas et al., 2017).

Moreover, several members of the OGDD family catalyze epigenetic modifications, e.g., demethylation of histones (Matilainen et al., 2017). OGDD enzymes referred to as TETs can convert 5-methylcytosine to 5-hydroxymethylcytosine (5hmC), thereby mediating DNA ten-eleven translocation (Spruijt et al., 2013). Indeed, hydroxymethylation of cytosine in DNA has unique roles to play in normal development and physiology, as well as in pathological processes, especially in organs containing terminally differentiated cells (Globisch et al., 2010). In connection with maladaptive remodeling of the heart, the state of 5-methylcytosine hydroxylation, particularly in genes related to energy production and mitochondrial function, undergoes pronounced alterations (Greco et al., 2016). The activities of various TET enzymes are largely dependent on access to their co-factor 2OG and its oxidized enantiomers L/D 2-hydroxyglutarate (2HG). L2HG is produced during adaptation to hypoxic and oxidative stress and this metabolite and the changes in 5hmC it causes may be important role in connection with maladaptive cardiac remodeling and heart failure.

The current investigation was designed to test our hypothesis that IDH2 is a major regulator of antioxidant defenses in the heart. We have elucidated the molecular and regulatory epigenetic functions of IDH2, 2OG and L2HG, as well as provided novel insights concerning the impact of antioxidative capacity on heart failure. These findings deepen our molecular understanding of redox homeostasis in cardiac tissue and may aid in the development of new approaches to the treatment of heart failure.

## Results

### Expression of IDH2 is downregulated in association with eccentric hypertrophic dilated cardiomyopathy

Previously, we performed RNA sequencing on left ventricular tissue (LV) from end-stage patients with dilated cardiomyopathy harboring mutations in lamin A/C (*LMNA*), RNA binding motif protein 20 (*RBM20*) and titin (*TTN*) (Sielemann et al., 2020). Here, to examine the involvement of mitochondria and related molecular pathways in heart failure, we performed Ingenuity Pathway Analysis (IPA) on these same data. This approach revealed significant enrichment in the levels of RNAs encoding proteins involved in pathways related to altered metabolic and redox homeostasis and dysfunction in mitochondria (**Figure 1A**). Subsequently, we confirmed these findings by performing IPA on published transcriptomic data from 64 samples of cardiac tissue from patients with dilated cardiomyopathy (van Heesch et al., 2019) (**Figure S1A**).

**Figure 1:**
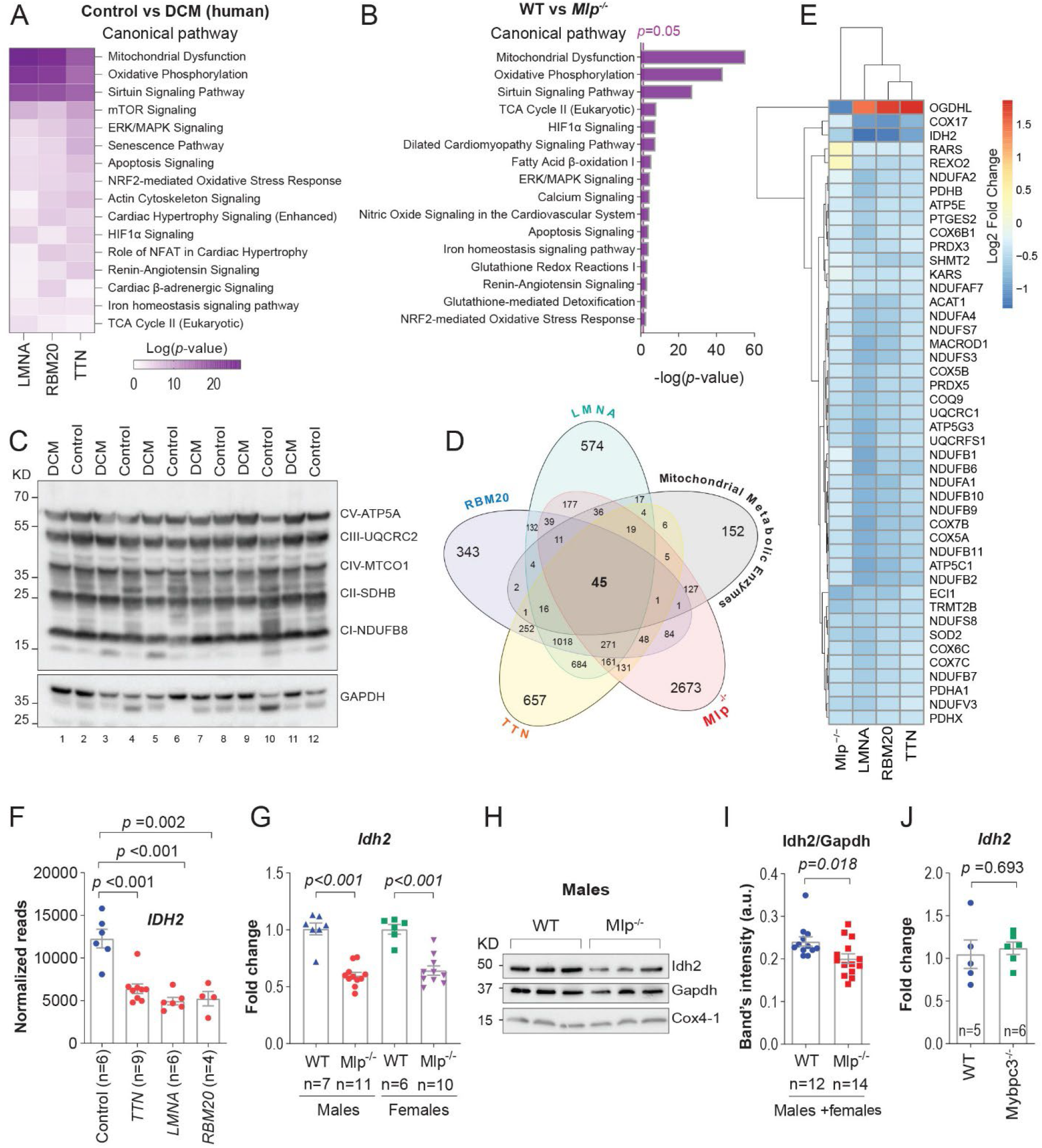
Mitochondrial dysfunction and downregulation of IDH2 expression in association with eccentric DCM. **A)** Selected pathways from IPA enrichment analysis on published transcriptomic data from left ventricle (LV) myocardium of patients with DCM (Sielemann et al., 2020). **B)** Selected pathways from IPA enrichment analysis on transcriptomic data from the LV of *Mlp*^*-/-*^ (males + females). **C)** Western blotting of OXPHOS complexes in the LV of patients with eccentric DCM (Samples’ metadata is described in table S1). **D)** Venn diagram of intersected genes encoding for mitochondrial metabolic enzymes, and differentially expressed genes (DEGs) in the LV of patients with DCM (described in figure 1A) or *Mlp*^*-/-*^ mice. Only DEGs with *p*<0.05 and adjusted *P* value < 0.01 were included in this analysis. **E)** A list over 45 genes encoding for mitochondrial metabolic enzymes that were commonly dysregulated in all data sets analyzed in figure (**1D**). **F)** *IDH2* expression inferred from RNA sequencing on the LV of patients with DCM described in figure 1A. **G)** qPCR analysis of *Idh2* expression in the LV of *Mlp*^*-/-*^. **H)** Western blotting of Idh2 in the LV of *Mlp*^*-/-*^. **I)** Quantification of the Western blotting of Idh2 in the LV of *Mlp*^*-/-*^ displayed in supplementary figure (**S1J**), which are from another larger set of males and females than the one displayed in figure (**1H**). **J)** qPCR analysis of *Idh2* expression in the LV of *Mybpc3*^*-/-*^ males. Bars are the mean values ±SEM with unpaired two-tailed *t*-test.

To unravel the underlying cause of this mitochondrial dysfunction, we subsequently analysed LV samples from patients with DCM, as well as from *Mlp*-deficient (*Mlp*^-/-^ or *Csrp3*^-/-^) mice, which develop an eccentric hypertrophy comparable to human DCM (Arber et al., 1997; Knöll et al., 2010). As in the human material, RNAs encoding proteins involved in pathways related to mitochondrial dysfunction were enriched in the *Mlp*^-/-^ mice as well (**Figures 1B and S1B**).

Transmission electron microscopy of the murine samples revealed no relevant alterations in mitochondrial numbers or structure (**Figure S1C and S1D**); nor were any alterations in the levels of components of complexes of the mitochondrial electron transport chain observed by Western blotting of either the human (**Figure 1C**) or murine samples (**Figures S1E and S1F)**. In agreement with these findings, respiration by mitochondria isolated from the transgenic animals was comparable to the corresponding findings with control mice (**Figure S1G**). Furthermore, no general common dysregulation in the expression of genes involved in the regulation of mitochondrial biogenesis, such as *PGC-1α* and *TFAM*, were observed between human and murine samples (**Figure S1H**). Thus, although the elevated pre- and afterload associated with heart failure demands more energy (Münzel et al., 2015), mitochondrial morphology and function appear to remain largely unchanged.

Since the energetic deficit associated with heart failure is, at least in part, related to alterations in substrate utilization (Munzel et al., 2017), we next identified genes encoding mitochondrial metabolic enzymes by intersecting the Mammalian Metabolic Enzyme and MitoCarta2 databases (Calvo et al., 2016; Corcoran et al., 2017). Of the 447 genes we found, 45 were significantly dysregulated in both patients with DCM and *Mlp*^-/-^ mice (**Figures 1D and 1E**). Intriguingly, *IDH2* was among the most significantly downregulated (**Figures 1F, 1G, and S1I**). The level of IDH2 protein in failing ischemic human hearts has been reported to be reduced (Ali et al., 2020), an observation which we confirmed in *Mlp*^-/-^ cardiac material, although this reduction was less pronounced than the decrease in the corresponding level of mRNA (**Figures 1H, 1I, S1J, and S1K**).

In contrast to *Mlp*^-/-^ mice, which display eccentric hypertrophy, the expression of *Idh2* was not dysregulated in the murine *Mybpc3*^-/-^ model of hypertrophic cardiomyopathy (HCM) (Carrier et al., 2004), where the animals demonstrate concentric hypertrophy (**Figure 1J**). Thus, downregulation of IDH2 appears to be characteristic specifically of DCM with eccentric hypertrophy.

### Oxidative stress downregulates *Idh2*

As mentioned above, in cardiac myocytes, IDH2 is primarily responsible for the regeneration of the mitochondrial NADPH required for enzymatic elimination of H_2_O_2_ (Nickel et al., 2015; Wagner et al., 2020). When we explored potential dysregulation of the expression of other genes involved in antioxidation in *Mlp*^-/-^ hearts, we observed upregulation of the *Nqo1* gene in both male and female mice, while the *Hmox1* gene was upregulated only in the male animals (**Figures 2A-C**).

**Figure 2:**
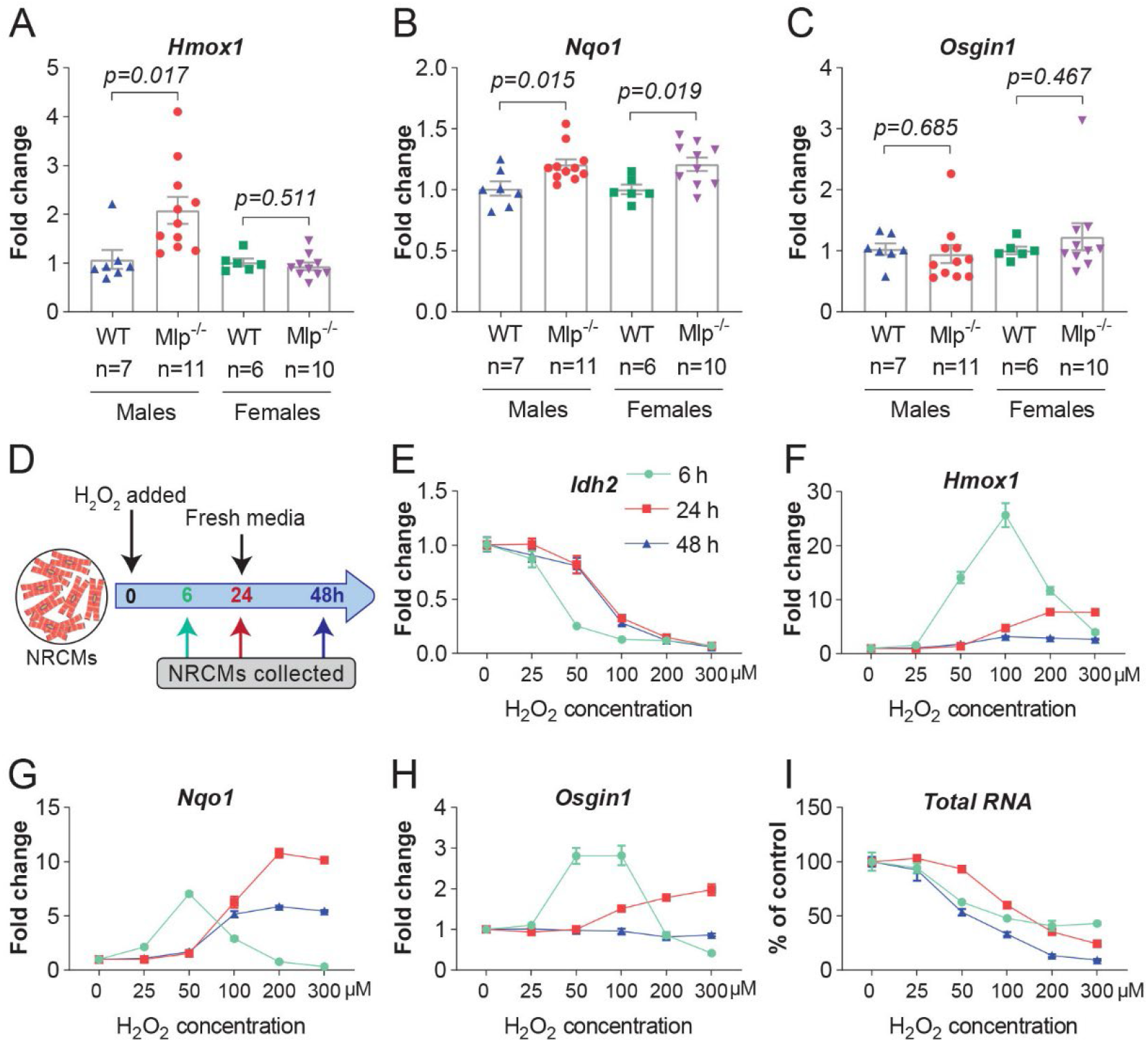
Oxidative stress downregulates *Idh2*. **A-C)** qPCR analysis of *Hmox1, Nqo1 & Osgin1* expression in LV myocardium of *Mlp*^*-/-*^, values are presented as the mean values ±SEM with two-tailed unpaired *t*-test. **D)** An experimental scheme depicting the treatment of NRCMs with increasing doses of H_2_O_2_. **E-H)** qPCR analysis of *Idh2, Hmox1, Nqo1* &*Osgin1* expression in H_2_O_2_-treated NRCMs normalized to untreated control. Equal amounts (5 ng) of total RNA were utilized for all qPCR reactions. **I)** Recovered amounts of extracted RNA from each well of H_2_O_2_-treated NRCMs. Values in **E-I** are the average of 4 individual wells ±SEM.

Next, when neonatal rat cardiomyocytes (NRCMs) were subjected to oxidative stress through exposure to increasing doses (0-300 μM) of H_2_O_2_ for 6 hours (**Figure 2D**), dose-dependent downregulation of *Idh2* occurred (**Figure 2E)**. Moreover, as the level of H_2_O_2_ gradually declined, this expression returned almost to normal, particularly in the case of cells exposed to low doses.

At the same time, further incubation of these cells with fresh medium free from H_2_O_2_ for an additional 24 hours did not raise the level of *Idh2* expression any further, indicating the potential presence of long-lasting effects (**Figure 2E**).

Utilizing RT-qPCR, we could confirm that exposure of NRCMs to oxidative stress induced the expression of *Hmox1, Nqo1* and *Osgin1*, downstream target genes for Nrf2 (**Figure 2F-H**). This effect was also associated with a loss of cardiomyocytes, as indicated by a decrease in the total amount of RNA recovered (**Figure 2I**). (This loss of RNA exerted no impact on the downregulation of Idh2 observed, since 5 ng total RNA was employed in all qPCR reactions.)

### Downregulation of Idh2 is associated with enhancement of its activity

To further examine the downregulation of IDH2 in response to oxidative stress, we next measured the activity of the Idh2 protein in subsarcolemmal mitochondria isolated from *Mlp*^-/-^ LV. Intriguingly, despite the lower levels of mRNA and protein (**Figures 1G-I**), this activity was higher in *Mlp*^-/-^ mice than in their WT littermates (**Figures 3A and 3B**). Moreover, this elevation was associated with higher levels of 2OG (**Figure 3C**).

**Figure 3:**
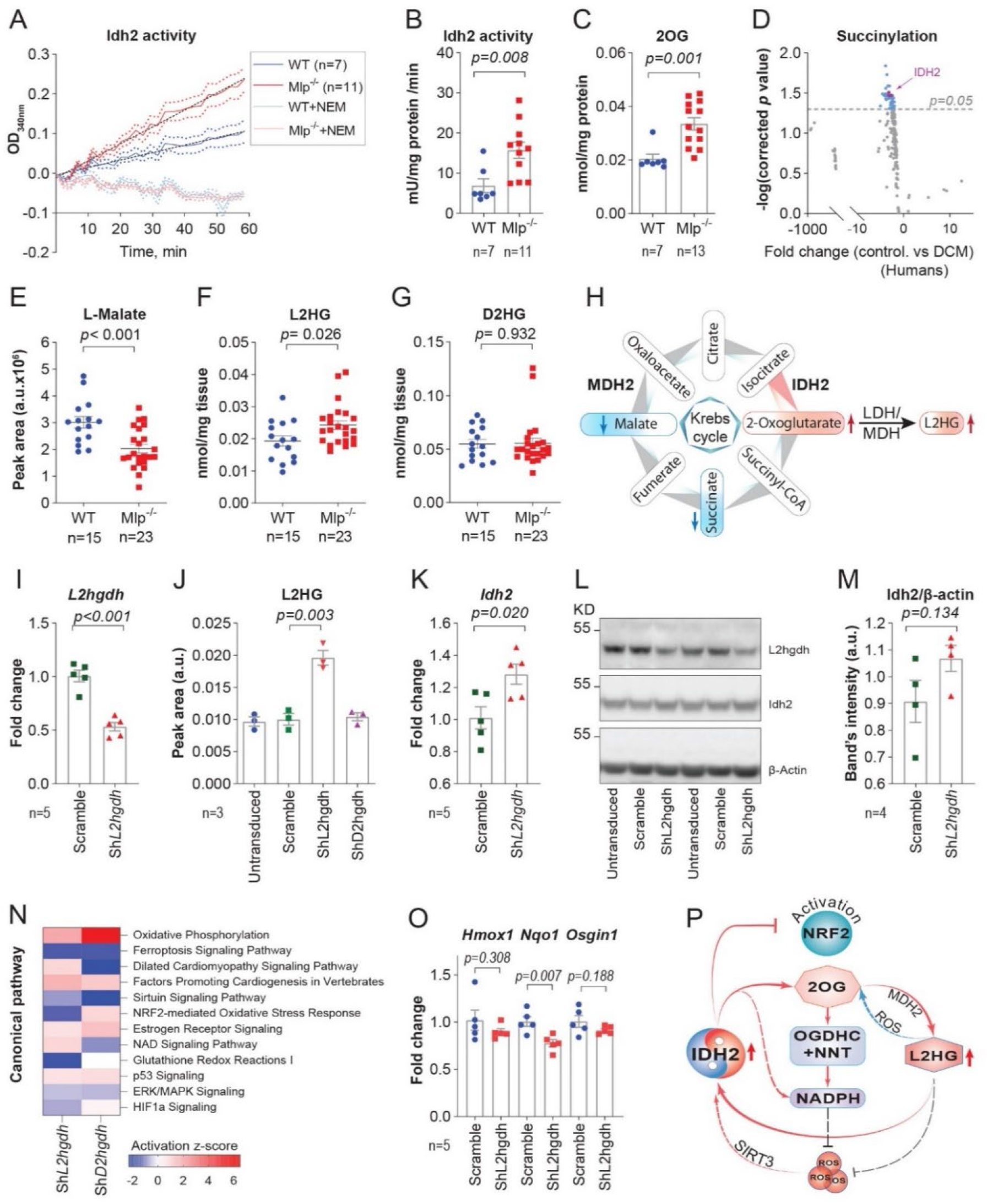
Antioxidative role of L2HG in the heart. **A and B)** Idh2 activity in cardiac mitochondria isolated from frozen LV myocardium of male *Mlp*^*-/-*^ with or without the Idh2-activity-inhibitor N Ethylmaleimide (NEM). **C)** 2OG levels in the LV of male *Mlp*^*-/-*^. **D)** Volcano plot for differentially succinylated peptides of mitochondrial metabolic enzymes in publicly available mass spectroscopic data on the LV of patients with ischemic cardiomyopathy (Ali et al., 2020). **E-G)** Targeted LC-MS analysis of L-malate, L2HG and D2HG levels in the LV of male and female *Mlp*^*-/-*^, normalized to tissue weight. **H)** A scheme depicting the metabolic remodeling of the Krebs cycle that underlays L2HG production **I-M)** NRCMs transduced with ShRNA targeting L2hgdh were analyzed for *L2hgdh* expression by qPCR in **(I)**; for L2HG levels by a targeted LC-MS in **(J)**; and for Idh2 expression on the mRNA level by qPCR in (**K**) and on the protein level by western blotting (in **L** and quantified in **M**). **N)** Selected pathways from IPA comparison analysis on transcriptomic data from NRCMs transduced with ShRNA targeting *L2hgdh* or *D2hgdh*. **O)** qPCR analysis of *Hmox1, Nqo1* and *Osgin1* expression in NRCMs transduced with ShRNA targeting *L2hgdh*.**P)** Proposed molecular mechanism for the antioxidative role of L2HG in the heart. All values are presented with the of mean value ±SEM, with unpaired two tailed *t*-test. Values in figures **I-O** are the average of n individual wells, whereby n number is indicated on the figures.

Among the post-translational modifications that modulate the stability and activity of IDH2, desuccinylation in response to oxidative stimuli elevates the activity of this enzyme (Zhou et al., 2016). Our analysis of publicly available mass spectroscopic data on the LV of patients with ischemic cardiomyopathy (Ali et al., 2020) revealed desuccinylation of multiple lysine residues in IDH2 (**Figure 3D**). Furthermore, the overall level of protein succinylation, as quantified by Western blotting, in *Mlp*^-/-^ hearts was lowered (**Figures S3A and S3B**), reminiscent of what has been observed in failing human hearts (Ali et al., 2020).

### The accumulation of L2-hydroxyglutarate associated with heart failure upregulates Idh2 and deactivates Nrf2

Under hypoxic conditions cellular levels of 2OG and lactate rise, while the level of L-malate falls, causing lactate (LDH) and malate dehydrogenases (MDH) to reduce 2OG to L2-hydroxyglutarate (L2HG). Subsequently, the L2HG formed participates in a redox couple system (2OG/L2HG) that regenerates NADH (Intlekofer et al., 2017; Oldham et al., 2015; Wise et al., 2011). Therefore, to characterize the metabolic remodeling associated with heart failure (Munzel et al., 2017), we applied targeted mass spectrometry.

In *Mlp*^-/-^ hearts the levels of L-malate and succinate were reduced (**Figures 3E and S3C**), while the level of lactate was elevated (**Figure S3D**). Chemical derivatization for more detailed analysis revealed that the level of L2HG, but not D2HG was enhanced (**Figures 3F-H**). This enhancement was not associated with upregulation of *Slc2a1*, a target gene for hypoxia inducible factor 1 (Hif1α), but was accompanied by induction of *Serpine1*, which is regulated by Hif2α (**Figure S3E**).

To examine the impact of L2HG and D2HG on the redox status of cardiomyocytes, the levels of these compounds in NRCMs were increased by ShRNA knockdown of *L2hgdh* and *D2hgdh*, i.e., the genes that encode mitochondrial dehydrogenases that oxidize L2HG and D2HG to 2OG, respectively. Approximately 50% knockdown of *L2hgdh* resulted in a two-fold increase in the level of L2HG (**Figures 3I and 3J**), while approximately 70% knockdown of *D2hgdh* elevated the level of D2HG 1.5-fold (**Figures S3F and S3G**). These findings are in line with previous reports that D2HG is elevated only in cancer cells carrying neomorphic mutations in IDH2 (Intlekofer et al., 2015; Oldham et al., 2015).

Although elevated levels of D2HG have been reported to downregulate IDH2 in a variety of cell types (Lin et al., 2015), we observed that elevated levels of L2HG in NRCMs were associated with upregulation of both Idh2 mRNA and protein (**Figures 3K-3M and S3H**), suggesting that cellular redox homeostasis was altered. Thereafter, transcriptomic profiling followed by IPA analysis revealed that while both L2HG and D2HG activate oxidative phosphorylation, only L2HG attenuates NRF2-mediated responses to oxidative stress responses, as well as glutathione redox reaction I (**Figures 3N and S3I**). We also confirmed deactivation of certain components of the Nrf2 pathway by L2HG through qPCR analysis of the expression of the downstream genes *Hmox1, Nqo1* and *Osgin1* (**Figure 3O**).

These data suggest that the increases in Idh2 activity and ensuing elevations in 2OG and L2HG levels enhance the antioxidative capacity of cardiomyocytes (**Figure 3P**). However, the functional reason for downregulation of *Idh2*, an essential component of cellular antioxidative defenses, in response to oxidative stress remained to be elucidated.

### Mutual redox regulation downregulates Idh2

In attempt to resolve the redox paradox described above, we treated NRCMs with 5 μM sulforaphane (SF) and/or 3 mM N-acetyl cysteine (NAC) and then monitored the expression of *Idh2* and three downstream target genes for Nrf2. SF enhances production of reactive oxygen species by mitochondria in cardiomyocytes (Rhoden et al., 2021); while NAC neutralizes free radicals, either directly through reaction with its sulfhydryl group and/or by increasing cellular levels of glutathione (Zafarullah et al., 2003).

SF induced the expression of the target genes while attenuating *Idh2* expression (**Figures 4B**), effects reminiscent of those produced by H_2_O_2_ (**Figures 2E-H**). NAC mitigated these effects of SF, while NAC alone upregulated *Idh2* and reduced the levels of downstream Nfr2 targets (**Figure 4B**), changes comparable to those that occur upon induction of L2HG (**Figure 3K**). These observations indicate the existence of an antioxidative control mechanism in which Idh2 and Nrf2 regulate each other’s activity (**Figure 4C**).

**Figure 4:**
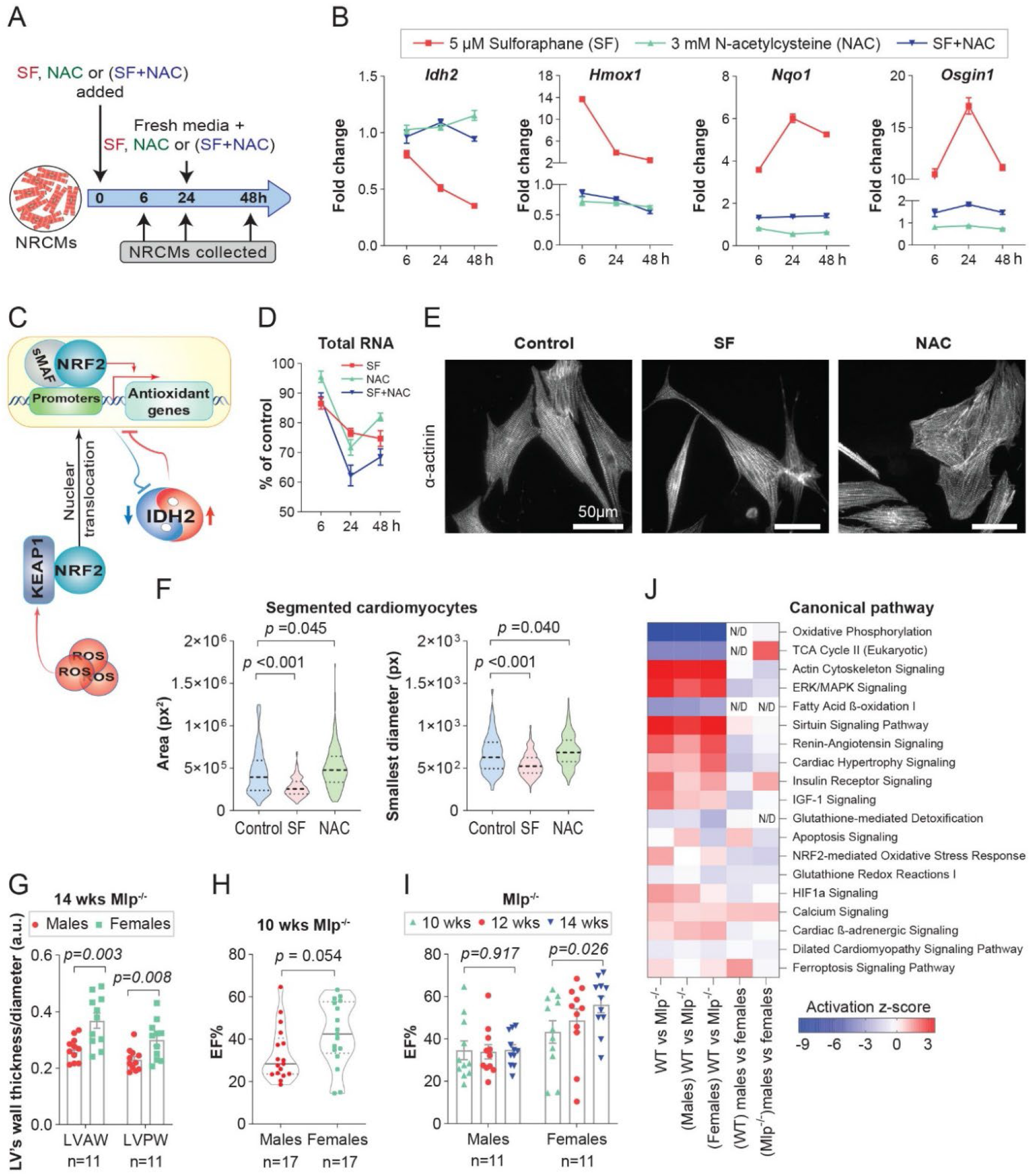
Morphology of cardiomyocytes and sex differences in responses to oxidative stress. **A)** An experimental scheme depicting the treatment of NRCMs with Sulforaphane (SF), N-acetyl cysteine (NAC) or in combinations. **B)** qPCR analysis of *Idh2, Hmox1, Nqo1* &*Osgin1* expression in treated NRCMs, normalized to untreated control. **C)** Proposed molecular mechanism for the Idh2-Nrf2 mutual regulation. **D)** Recovered amounts of total RNA extracted from the treated-NRCMs. Values in figures **B** and **D** are the average value of 4 individual wells ±SEM. **E)** NRCMs stained for α-actinin and imaged with a spinning disk confocal microscope to visualize the effect of SF and NAC treatments on cell shape and structure. **F)** ImageJ analysis of cell surface area and the minimum caliper diameter of the SF or NAC-treated NRCMs, measured in pixel (px). The median is shown on the plot, with the interquartile range and unpaired one-tailed *t*-test. **G)** LV anterior (LVAW) or posterior (LVPW) wall thickness, normalized to LV diameter, in male and female *Mlp*^*-/-*^ mice. The values are the average of systolic and diastolic ones. Bars are the mean values ±SEM with unpaired t-test. **H**) LVEF in male and female *Mlp*^*-/-*^ mice at the age of 10 weeks. The median is shown on the plot, with the interquartile range and unpaired t-test. **I)** Changes in LVEF of male and female *Mlp*^*-/-*^ mice over time. Bars are the mean values ±SEM with paired two-tailed *t*-test. **J)** Selected pathways from IPA comparison analysis on transcriptomic data from the LV of male and female *Mlp*^*-/-*^ mice.

These findings led us to hypothesize that the robust antioxidative capacity of the heart explains, at least in part, the failure of clinical attempts to treat heart failure by ameliorating oxidative stress. To test the potential consequences of these findings for treatment of patients with heart failure, we needed first to establish an appropriate model.

### Intracellular redox homeostasis modulates the morphology of cardiomyocytes

As reflected in the decrease in the total amount of RNA recovered, treatment of NRCMs with SF and/or NAC for 24 hours led to significant loss of cells. However, upon treatment with SF for an additional 24 h there was little further loss, while with NAC there was even a slight increase in the amount of RNA recovered (**Figure 4D**).

Cardiomyocytes do not divide and potential proliferation of contaminating cardiac fibroblasts was inhibited by the presence of horse serum in combination with cytarabine, an inhibitor of mitosis, in the culture medium. Therefore, we hypothesized that the blunting of cell death and the increase in RNA content observed upon prolonged treatment with NAC is the result not only of an altered antioxidant response, but also of remodeling of cardiomyocyte morphology. Indeed, SF was found to lead to sarcomeric disarray and cardiomyocyte atrophy, while NAC promoted cellular hypertrophy. (**Figures 4E, 4F and S4A**). These findings indicate that intracellular redox homeostasis is involved in regulating the morphology of cardiomyocytes.

### Sex differences in responses to oxidative stress

Since the hearts of male *Mlp*^-/-^ mice experienced more oxidative stress than those of female animals (**Figure 2A**), we wondered whether this difference would be reflected in the properties of the heart. These mice develop DCM at a very early age, but with extensive variability in the disease severity (Arber et al., 1997). Echocardiograpy revealed that the walls of the left ventricle of 14-week-old male mice were less thick than those of females, but with no significant difference at 10 weeks of age (**Figure 4G and S4B**). Moreover, at the younger age females exhibited a greater fraction of LV ejection (LVEF) (**Figure 4H**), a difference that became more pronounced with age as the female LVEF improved, with no change in the males (**Figure 4I**). This improvement in the LVEF in female animals was associated with an increase in the thickness of the wall of the LV, with no such change in the case of the males (**Figure S4C**). In general, this remodeling of the LV is consistent with the effects of redox homeostasis on cardiomyocyte morphology described above.

Thus, the phenotype of the heart of male *Mlp*^-/-^ mice is more severely DCM than that of the females. In contrast, no such differences between male and female WT animals were observed. Moreover, two-way ANOVA analysis of certain cardiac parameters in mice of both genotypes indicated that the effect of genotype on the thickness of the LV wall was dependent on sex (**Figure S4D**).

To characterize oxidative stress in male and female *Mlp*^-/-^ hearts in greater detail, we carried out transcriptomic profiling at 12 weeks of age, followed by comparative IPA analysis. This showed that the female animals exhibited less extensive changes in Nrf2-mediated responses to oxidative stress and in glutathione redox reaction I (**Figure 4J**), reflecting, as expected, their lower level of oxidative stress. Altogether, these results indicate that there is a correlation between the level of oxidative stress and severity of the phenotype.

### Activation of Nrf2 improves cardiac function in male, but not female *Mlp*^-/-^ mice

In light of the considerable defensive antioxidative capacity of the heart, we hypothesized that patients with heart failure involving impairment of this capacity would benefit more from redox treatments. To test this hypothesis, we exploited the more adequate defenses against oxidative stress demonstrated by female than male *Mlp*^-/-^ mice.

To this end, we activated Nrf2-mediated antioxidant responses by destabilizing the Keap1-Nrf2-Cul3 complex, which ubiquitinylates Nfr2, by treatment with the small novel AZ925 molecule, which blocks the BTB domain of Keap1. Such destabilization releases free Nrf2 into the cytoplasm, which then translocates into the nucleus and activates transcription of antioxidant genes such as *Hmox1, Noq1* and *Osgin1*.

Consistent with our hypothesis, daily gavage with AZ925 for 30 days and monitoring with echocardiography on days 0, 14 and 30 (**Figure 5A**) revealed significant improvement of LVEF in male, but not in female *Mlp*^-/-^ mice (**Figures 5B-E**). Moreover, this improvement in the male animals was associated with an increase in the thickness of the left anterior wall during systole, whereas no such change occurred in the females (**Figure S5A and S5B**).

**Figure 5:**
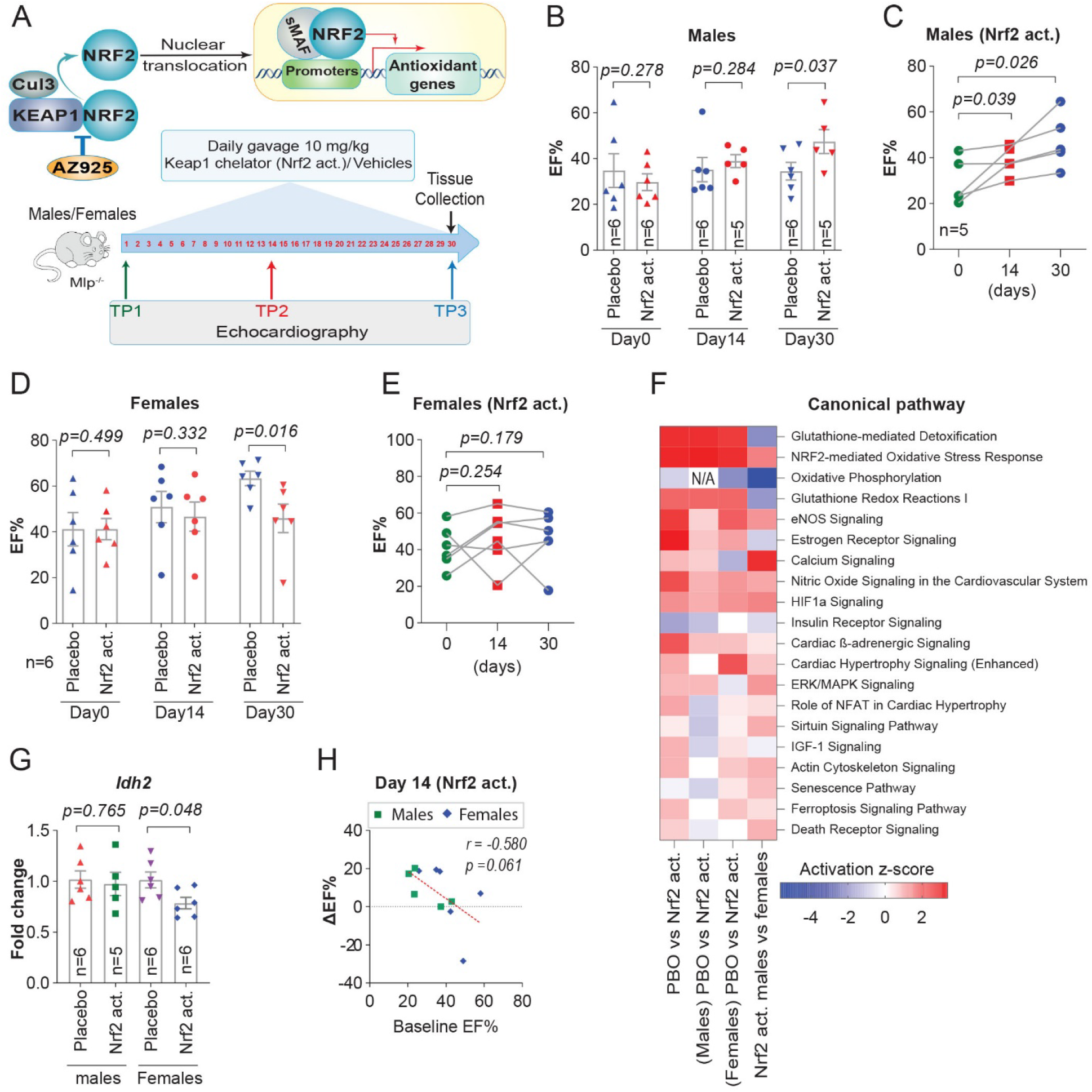
Activation of Nrf2 improves cardiac function in male, but not female *Mlp*^*-/-*^ mice. A scheme for the experimental settings of the in-vivo Keap1 chelator (Nrf2 activator) study and for underlying molecular mechanism. **B-E)** Ejection fraction (EF%) of male (**B**) and female (**D**) *Mlp*^*-/-*^ mice treated with Nrf2 activator or with the vehicle control. The changes over time in EF% for individual animals treated with the Nrf2 activator are shown for males in (**C**) and for females in (**E**). Bars are the mean values ±SEM with unpaired one-tailed *t*-test (in **B** and **D**) or paired one-tailed *t*-test (in **C** and **E**). **F)** Selected pathways from IPA comparison analysis on transcriptomic data from the LV of *Mlp*^*-/-*^ treated with Nrf2 activator or with the vehicle control. **G)** qPCR analysis of *Idh2* expression in the LV of *Mlp*^*-/-*^ treated with Nrf2 activator, normalized to the vehicle control group. Values are the mean fold change ± SEM with unpaired two-tail *t*-test. **H)** Spearman’s correlation between the baseline EF% of male and female *Mlp*^*-/-*^ mice and the ΔEF% after 14 days of treatment with Nrf2 activator.

Moreover, as determined by RNA sequencing followed by IPA analysis, this treatment with AZ925 potently activated pathways involved in antioxidant responses -- including glutathione-mediated detoxification, the Nrf2-mediated response to oxidative stress, and glutathione redox reactions – in mice of both sexes (**Figure 5F**). In addition, induction of *Hmox1, Noq1* and *Osgin1*, genes encoding downstream targets of Nfr2, in the treated animals was confirmed (**Figure S5C-E**).

Interestingly, treated female *Mlp*^-/-^ mice exhibited more potent activation of the Nrf2-mediated responses to oxidative stress, but weaker activation of glutathione-mediated pathways. These differences are consistent with our observation that untreated *Mlp*^-/-^ females demonstrate less oxidative stress than males (**Figure 4J**), as well as with the pronounced additional downregulation of Idh2 in treated female, but not male mice (**Figure 5G**).

Altogether, these findings indicate that patients with heart failure and insufficient cardiac antioxidant capacity would, indeed, benefit more from antioxidant treatment to a degree dependent on LV systolic function and the level of oxidative stress. However, this certainly does not imply that female patients with heart failure would not benefit from such treatment. For example, such treatment initially improved the cardiac phenotype of female *Mlp*^-/-^ mice with a low ejection fraction (EF<40%) (**Figure 5E**). Moreover, there was a general negative correlation (*r* = -0.580) between baseline LVEF and improvement in this parameter following 14 days of treatment for mice of both sexes (**Figure 5H**).

### Unique epigenetic control of Idh2 expression and redox response

The experiments described above reveal that expression of *Idh2* is regulated through the antioxidative capacity. Since the 2OG/L2HG ratio is a potent modulator of the activities of Tet1-3 enzymes (Xu, Yang et al. 2011), we examined the potential involvement of epigenetic changes in this context. Although D2HG appears to be involved in remodeling of the cardiac epigenome and consequent mitochondrial dysfunction (Akbay, Moslehi et al. 2014, Karlstaedt, Zhang et al. 2016), no such role for L2HG has yet been reported.

Accordingly, we assessed the level and distribution of 5-methylcytosine (5mC) and 5-hydroxymethylcytosine (5hmC) in the DNA of *Mlp*^-/-^ hearts by developing a tool to analyze a publicly available whole-genome oxidative bisulfite (OxBS) and bisulfite (BS) sequencing data with single-base pair resolution on *Mlp*^-/-^ hearts (SRA accession ID: PRJNA328890). Our analysis indicated that the average methylation percentages of the total CpGs in the murine genome were 58.7% for 5mC and 4.4% for 5hmC. In *Mlp*^-/-^ hearts, there were trends toward increased levels of 5mC and fewer 5hmC (**Figure 6A and 6B**). Moreover, principal component analysis demonstrated that 5mC and 5hmC are distributed differently in the cardiac genomes of the transgenic and wild-type animals (**Figure 6C and 6D**). At the same time, there was no significant difference in either the level of mRNA encoding Tet1-3 or its enzymatic activity (**Figures S6A and S6B**). These results indicate that in cardiomyocytes cofactors of Tet1-3 enzymes, including L2HG, may modulate their activities, thereby altering the genomic profiles of 5mC and 5hmC in a manner that contributes to the development of heart failure.

**Figure 6:**
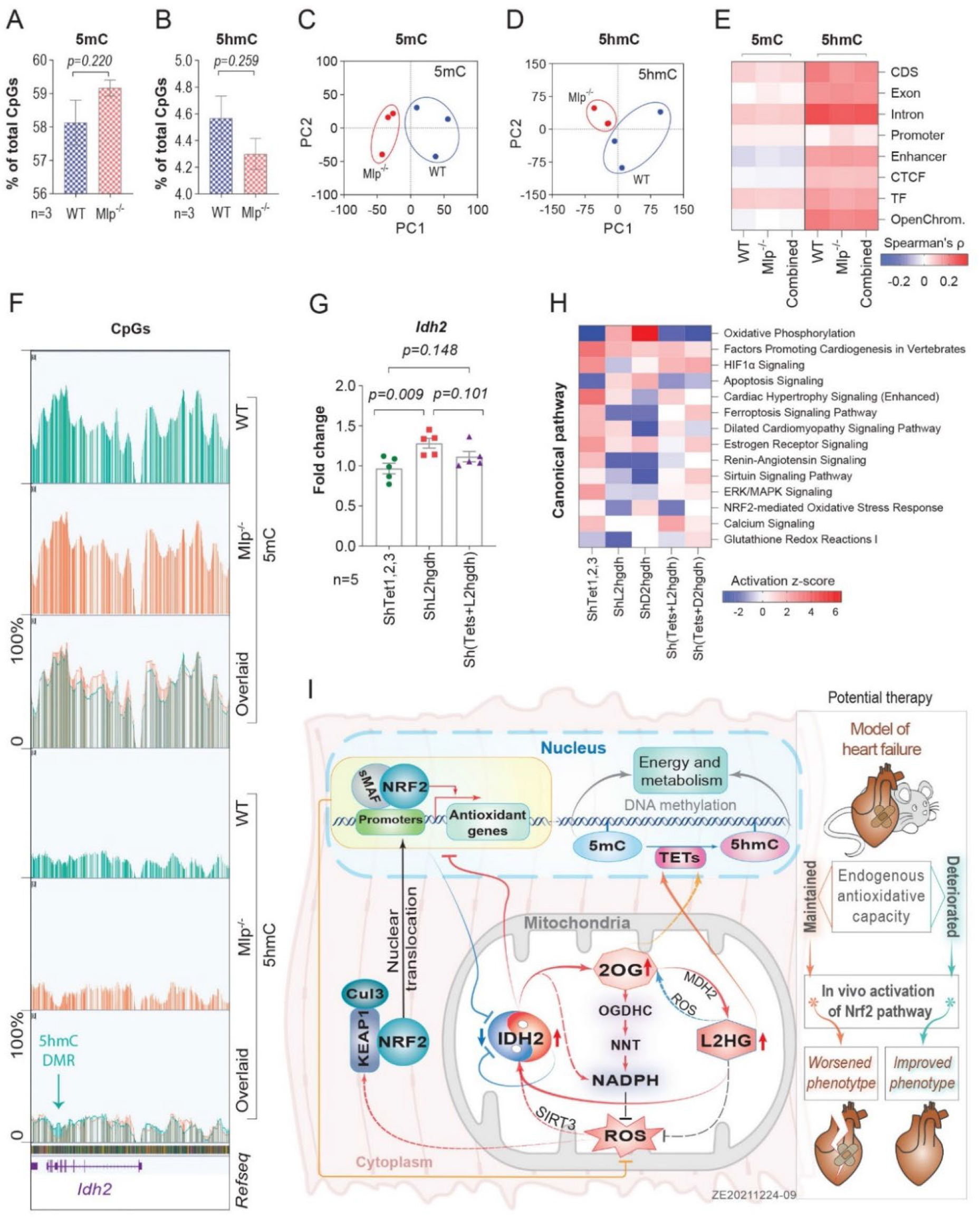
Unique epigenetic control of Idh2 expression and redox response. **A and B)** Percentages of whole genome 5mC or 5hmC-methylated CpGs in the myocardium of *Mlp*^*-/-*^ mice **C and D)** Principal component analysis on the distribution of whole genome 5mC and 5hmC across biological samples (n=3). **E)** Spearman correlation between the expression of individual genes and the proportion of methylation in each functional region of individual samples. (CDS: Coding regions, CTCF: Transcriptional repressor CTCF binding sites, TF: transcription factors binding sites.). The values presented are the average of three biological replicates. **F)** A screenshot from the Integrative Genomic Viewer for actual methylation levels in *Idh2* locus in chromosome 7 of the murine genome. **G)** qPCR analysis of *Idh2* expression in NRCMs transduced with ShRNA targeting *Tet1-3* and/or *L2hgdh* normalized to their respective scramble control. **H)** Selected pathways from IPA comparison analysis on transcriptomic data of NRCMs transduced with ShRNA targeting *Tet1-3, L2hgdh, D2hgdh*, or *Tet1-3* combined with *L2hgdh* or *D2hgdh*. Bars in figures **A, B** and **G** are the mean values ± SEM with unpaired two-tailed *t*-test. **I)** A summary over the whole proposed molecular mechanism in this work.

Functional analysis of the distributions of 5mC and 5hmC in the cardiac genomes of *Mlp*^-/-^ and WT mice revealed that *Mlp*^-/-^ hearts contained lower levels of 5hmC in all functional regions, with no differences in 5mC levels (**Figures S6C**). Subsequent, Spearman’s rank correlation between the levels of expression of individual genes and the proportion of methylation in each functional region showed a noteworthy positive correlation between gene expression and intronic levels of 5hmC (**Figure 6E**). Moreover, computing differentially methylated regions (DMRs) indicated that introns, followed thereafter by promoters harbored the highest number of the identified DMRs (**Figure S6D**). In addition, the genes that contained DMRs in their introns encoded a particularly high number of proteins involved in hypertrophic remodeling and regulation of redox homeostasis (**Figure S6E**).

The genes in *Mlp*^-/-^ hearts that contained lower levels of 5hmC in an intron included *Idh2* (**Figure 6F**), a difference that was associated with a lower level of the expression of this gene (**Figure 1G**). To examine whether the levels of Tet1-3 enzymes influence *Idh2* expression, we knocked down *Tet1-3* in NRCMs with ShRNA, but this only reduced the levels of *Tet1* and *Tet3* mRNA (**Figure S6F**). At the same time, simultaneous knockdown of Tet1/3 and L2hgdh attenuated the induction of *Idh2* caused by elevated levels of L2HG (**Figure 6G**), indicating that 5hmC plays a role in mediating this induction.

Transcriptomic profiling revealed that knockdown of Tet1/3 influenced a number of pathways involved in regulating oxidative stress and mitochondrial function (**Figure S6G**). Moreover, simultaneous knockdown of Tet1/3 and L2hgdh or D2hgdh affected the enrichment in many metabolic pathways mediated by L2HG and D2HG that we observed previously (**Figure 6H**). These findings constitute evidence that Tet1/3 and the 5hmC it generates are involved in regulating energy metabolism and mitochondrial function in cardiomyocytes (**Figure 6I**).

## Discussion

The present comprehensive approach – involving a combination of metabolic, transcriptional and epigenetic data -- reveals unique features of cardiac antioxidant defenses. For instance, we show here that IDH2 governs robust antioxidative defenses in mitochondria and coordinates these with other defensive pathways (**Figure 6I**), which probably influences cardiomyocyte morphology. In addition, by activating Nrf2 in a murine model (*Mlp*^*-/-*^) of heart failure through treatment with a small therapeutic molecule, we show that boosting endogenous antioxidative capacity improves LV function, particularly when this capacity is inadequate. This finding has important consequences in connection with antioxidant treatment of patients with heart failure.

### In connection with heart failure the antioxidative defenses of cardiomyocytes are highly orchestrated

Here, we observed that both acute and chronic exposure to pro-oxidants led to dose- and time-dependent downregulation of *Idh2*, which may have appeared dysfunctional. However, we subsequently demonstrated that this downregulation of *IDH2* was actually associated with an increase in total IDH2 activity as a consequence of posttranslational modifications. Moreover, this increase in IDH2 activity elevated intracellular levels of 2OG and L2HG, which also mitigate oxidative damage (Oldham et al., 2015).

Furthermore, mitochondria in females have been reported to exhibit less oxidative damage and to contain higher levels of reducing equivalents than those of males (Borrás et al., 2003). In this context, we observed that activation of pathways by oxidative stress was less pronounced in the hearts of female than male *Mlp*^-/-^ mice. This may explain the additional downregulation of *Idh2* in female, but not male *Mlp*^-/-^ hearts upon activation of the Nrf2 pathway by AZ925.

Altogether, these observations indicate that the downregulation of *Idh2* in response to oxidative stress may reflect the presence of a negative feedback loop. Further evidence for this proposal was provided by our findings that *in vitro* activation of the Nrf2 pathway with prooxidants downregulated *Idh2*, whereas deactivation of this same pathway by NAC or L2HG upregulated *Idh2*.

In this context, NRF2 has not been reported to be an upstream regulator of IDH2, whose gene lacks the antioxidant response element (ARE) in its promoter. This suggests that IDH2 and NRF2 are intimately interconnected in some other manner. Both *Idh2*^-/-^ and *Nrf2*^-/-^ strains of mice are viable, but with reduced defenses against oxidative stress and enhanced susceptibility to heart failure (Ku et al., 2015; Li et al., 2009). Many of the genes involved in antioxidant defenses mediated by glutathione are induced by NRF2 (Thimmulappa et al., 2002) and in cardiomyocytes the NADPH required to regenerate the reduced form of glutathione (GSH) is provided primarily by IDH2 and downstream OGDH coupling to NNT in mitochondria (Nickel et al., 2015; Wagner et al., 2020). These observations indicate that the cellular antioxidative machinery is closely orchestrated to mount effective defenses without overwhelming the cell with antioxidant equivalents.

### IDH2, 2OG and L2HG form a novel antioxidative feedforward cycle

The elevations in the levels of 2OG and L2HG in response to oxidative stress indicate that they may be involved in maintaining cellular antioxidative capacity. Indeed, 2OG enhances NADPH regeneration by cardiac mitochondria (Wagner et al., 2020) and elevates L2HG, thereby producing an L2HG/2OG redox-couple (Oldham et al., 2015). In addition, we show here that L2HG upregulates Idh2 and activates antioxidative responses. Altogether, these observations indicate that IDH2, 2OG and L2HG participate in an antioxidative feedforward loop that not only prevents oxidative damage, but also senses and regulates the antioxidative status of mitochondria. Consequently, the pronounced downregulation of *IDH2* in eccentric DCM hearts may reflect extensive activity of this cycle.

All components of this cycle are located in the mitochondria, where major formation of ROS occurs (Ide et al., 1999). This observation indicates that this antioxidative defense is not restricted to the heart, but also plays an important role in other metabolically active organs that express NNT, such as the liver, kidney, and spleen. However, the details concerning this feedforward cycle and its significance in other organs remain to be explored experimentally.

### 5hmC regulates IDH2 and the response to oxidative stress

Perhaps the most striking finding presented here is the profound impact of intronic 5hmC on gene expression. In a murine model of heart failure due to transverse aortic constriction, loss of 5hmC was recently reported to be associated with lowered expression of genes involved primarily in regulating energy and metabolism (Greco et al., 2016). Our analysis of high-resolution OxBS whole-genome sequencing data enabled us to demonstrate more specifically that this effect is due to alterations in intronic 5hmC.

In addition, we observed that 5mC and 5hmC in other regions of genes exert an impact on their expression, highlighting the importance of such modifications for the affinity of transcriptional factors to regulatory sequences (Spruijt et al., 2013). Moreover, our knockdown of Tet1/3 provided direct evidence that activation of antioxidative pathways and the marked induction of Idh2 expression by L2HG are mediated, at least in part, through epigenetic modifications. Together, these findings constitute evidence that 5hmC, 5mC and L2HG play critical roles in epigenetic processes that link oxidative stress to mitochondrial function, with intronic 5hmC as a fundamental regulator of gene expression in the heart.

### Clinical importance, limitations, and future perspectives

On the basis of the results described here, we propose that IDH2 is a major regulator of mitochondrial antioxidant defenses in the heart. Accordingly, assessment of this antioxidant capacity may help predict the chances of success when subjecting cardiac oxidative stress or, perhaps, certain cancers as well to antioxidant treatment. In this context, application of such individualized treatment in the clinic would require noninvasive biomarkers.

As described here, L2HG and 2OG represent promising novel candidates for such biomarkers and, in fact, elevated levels of 2OG have been detected in the plasma of patients with DCM (Dunn et al., 2007). However, cellular antioxidative defenses are complex and there are almost certainly additional mechanisms that should be taken into consideration in this connection. Nonetheless, our current findings provide important novel insights concerning cellular redox biology that will hopefully aid in the development of more effective personalized treatments.

Moreover, we demonstrate here that during cardiac remodeling hydroxymethylation of cytosine in specific regions of genes is a potent regulatory of gene expression. Unfortunately, because of its extremely low levels in the genome, elucidating this regulatory role of 5hmC in greater detail would require high sequencing coverage (> 200X), which is enormously costly at present. Hopefully, emerging third-generation sequencing and other future advances will soon facilitate such investigations.

## Supporting information

Supplementary figures and a detailed description of the methods

## Acknowledgements

We acknowledge all people that have contributed to this study, either by providing funds or technical assistance. We want to particularly acknowledge Mattias Svensson at the department of medicine, KI, for his valuable intellectual input and feedback, Oscar Franzén for his valuable input in the epigenetic bioinformatic analysis, Byambajav Buyandelger for her valuable intellectual input and feedback, as well as for preparing samples for TEM. Anne-Laure Lainé, for preparing AZ925, David Brodin at the BEA facility, KI, Huddinge, for his support in analyzing the ShRNA-KD RNA seq data, Kjell Hultenby for performing the TEM and quantifying the images obtained, Amy Li for her valuable input with the metadata of human heart samples from the Sydney biobank, Sara Fernandez Leon and Charlotte Webster for their technical assistance, and Joseph W DePierre for the scientific English editing of the manuscript.

We would like to acknowledge the Single-cell core facility at the Flemingsberg campus (SICOF) of Karolinska Institute for their sequencing. This facility is supported by grants from the KI Department of Medicine (MedH) and KI Infrastructure, as well as the infrastructure for the Strategic Research Area (SFO) on Stem Cells and Regenerative Medicine.

The florescence microscopy was performed at the Live Cell imaging Core facility/Nikon Center of Excellence at Karolinska Institute, with support from the Swedish Research Council, KI infrastructure, and the Centre for Innovative Medicine.

We also acknowledge Gisele Miranda for her input in the analysis of fluorescence images. The BioImage Informatics Facility for carrying out image analysis is funded by SciLifeLab, the National Microscopy Infrastructure, NMI (VR-RFI 2019-00217), and the Chan-Zuckerberg Initiative. The Proteomics Biomedicum core facility of Karolinska Institutet for performing the mass spectrometric analysis.

Antibodies against VDAC, COX4I1, and NDUFB8 were a kind gift from Peter Rehling, Göttingen, and we thank Berkan Arslan for performing the Western blotting.

Christoph Maack is supported by the German Research Foundation (DFG; SFB 894, TRR-219; Ma 2528/7-1) and the German Ministry of Education and Research (BMBF, 01EO1504). Nikolay Oskolkov is supported financially by the Knut and Alice Wallenberg Foundation as part of the National Bioinformatics Infrastructure Sweden at SciLifeLab

## Author contributions

**Z.E**.: Conceived, designed, conducted, and supervised all experiments; led the analysis of the epigenetic data; analyzed and interpreted the data; and wrote the manuscript.

**M.B.H**.: Echocardiography; jointly contributed to the design and animal gavage in connection with the study involving the inhibitor of Keap1; assay of Tets activity; interpreted the data and edited the manuscript.

**H.S**.: Echocardiography and bioinformatic analysis of BS and OxBS data; differential gene expression analysis of all RNA seq data.

**N.O**.: Development and supervision of the procedures for analysis of epigenetic data.

**B. B**.: Preparation of samples for TEM.

**J. L**.: Preparation of DNA libraries and sequencing in connection with the in vivo Keap1 inhibitor study.

**F.K. & A.W**.: Alignment and demultiplexing of RNA seq data obtained in the Keap1 inhibitor study.

**C.D.R**.: Provision of the human heart samples from the Sydney biobank.

**D.K**.: IPA analysis of the human heart samples.

**R.J**., **R.B**., **T.J**., **E.F**.: Discovery of AZ925 and enabling the pilot study.

**A.N**., **M.K**., **J.D**., **C.M**.: Measurement of the rate of respiration by mitochondria isolated from *Mlp*^*-/-*^; Western blotting of VDAC, COX4I1, and NDUFB8; interpreted the data, and provided scientific input and editing of the manuscript.

**M.F, A.D. Cl.B:** Scientific contribution to the manuscript.

**L.H.L**.: Supervised and provided scientific and editing input to the manuscript.

**A.V**.: Design, acquisition, analysis and interpretation of all mass spectrometric data, edited the manuscript. **Ch.B**.: Supported the experiments, interpreted the data, provided scientific and editing input to the manuscript.

All authors contributed to the final editing of this manuscript.

### Lead Contact and Materials Availability

The supplementary file contains supplementary figures and a detailed description of the methods. Requests for further information and resources should be directed to the lead contact, Zaher ElBeck.

