## Supplementary figures and a detailed description of the methods for "Epigenetic modulators link mitochondrial redox homeostasis to cardiac function"

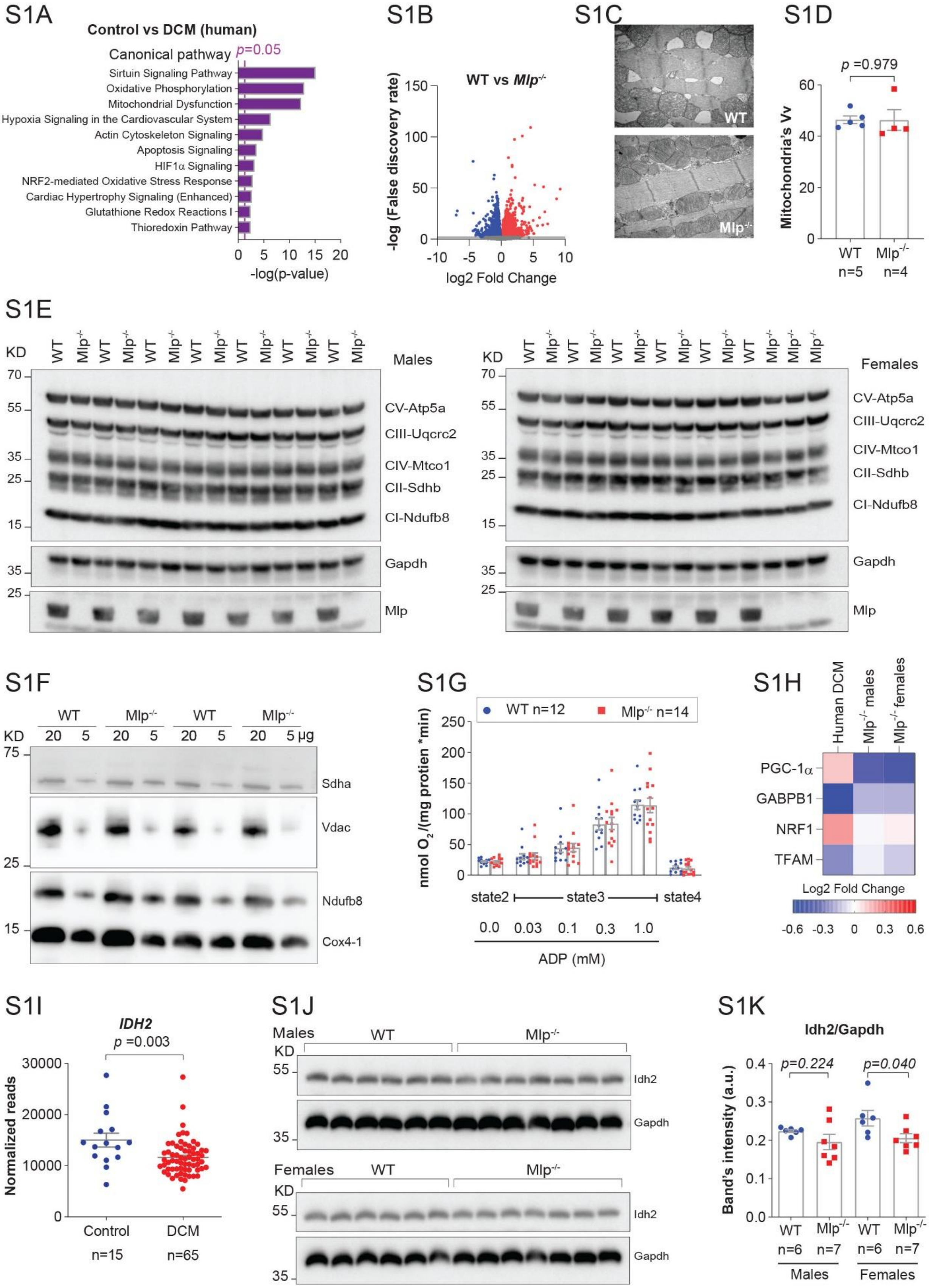

**Supplementary Figure S1: Mitochondrial dysfunction and downregulation of IDH2 expression in association with eccentric DCM**

**S1A)** Selected pathways from IPA enrichment analysis on published transcriptomic data from 64 myocardial samples from patients with DCM (van Heesch et al., 2019).

**S1B**) Volcano plot of transcriptomic data on the left ventricle (LV) myocardium of male and female *Mlp*$^{-/-}$ mice.

**S1C&D)** Images of LV myocardium of *Mlp*$^{-/-}$ mice, captured with transmission electron microscopy, and the subsequent quantification of mitochondrial volume density (Vv).

**S1E)** Western blotting of OXPHOS complexes in the LV of male (left panel) and female (right panel) *Mlp*$^{-/-}$ mice.

**S1F)** Western blotting of mitochondrial proteins (Vdac, Sdha, Ndufb8 & Cox4-1) in 5 and 20 µg of freshly isolated mitochondria from the LV of *Mlp*$^{-/-}$ mice.

**S1G)** Oxygen consumption in freshly isolated mitochondria from mitochondria from the LV of *Mlp*$^{-/-}$ mice.

**S1H)** Expression of genes involved in the regulation of mitochondrial biogenesis in the LV of patients with DCM (Described in figure S1A) and, male and female *Mlp*$^{-/-}$ mice, analyzed by RNA sequencing.

**S1I)** *IDH2* expression in the LV of patients with eccentric DCM described in figure S1A, analyzed by RNA sequencing.

**S1J &K)** Western blotting of Idh2 in the LV of male and female *Mlp*$^{-/-}$ mice and a subsequent quantification of the bands' intensity (The quantification shown in the main figure 1I is the merge of males and females in these two plots).

Bars are the mean value ± SEM with unpaired two-tailed t-test.

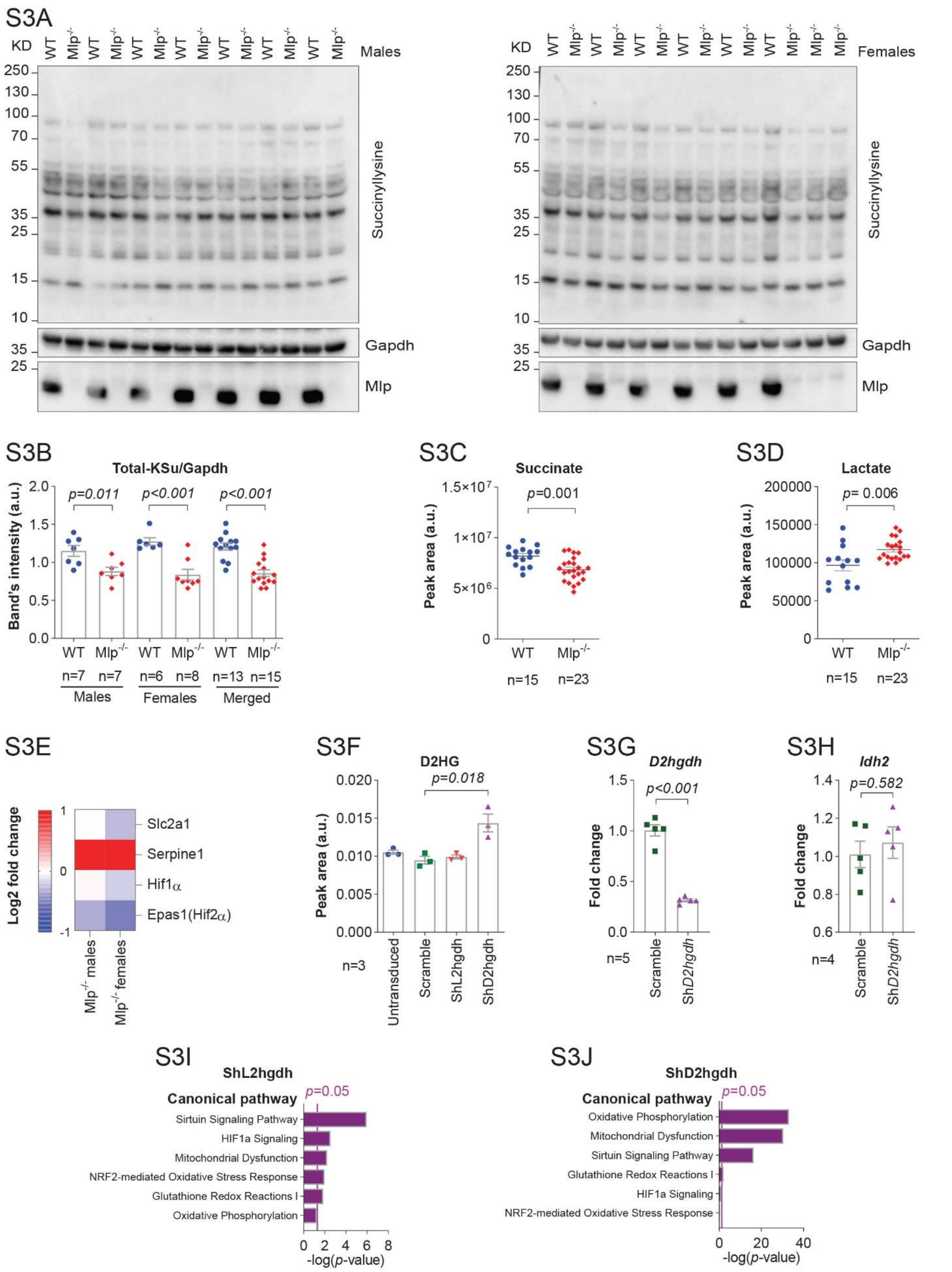

**Supplementary Figure S3: Antioxidative role of L2HG in the heart**

**S3A&B)** Western blotting of succinylated proteins in the LV of male and female *Mlp*$^{-/-}$ mice with total bands' intensity quantified in **(S3B)**. Bars are the mean values ±SEM with unpaired two-tailed *t*-test.

**S3C&D)** Targeted LC-MS analysis of succinate and lactate levels in the LV of *Mlp*$^{-/-}$ mice. Horizontal lines are the mean values ±SEM with unpaired two-tailed *t*-test.

**S3E)** Expression of genes regulating hypoxia in the LV of *Mlp*$^{-/-}$ mice analyzed by RNA sequencing.

**S3F-H)** Transduced NRCMs with ShRNA targeting *D2hgdh* were analyzed for the of D2HG by a targeted LC-MS (in **S3F)**; and for the expression of *D2hgdh* and *Idh2* by qPCR **(**in **S3G&H).** Values are the average of n individual wells ±SEM with unpaired two-tailed *t*-test, whereby n number is indicated on the figures.

**S3I&J)** Selected pathways from IPA enrichment analysis on transcriptomic data from NRCMs transduced with ShRNA targeting *L2hgdh* or *D2hgdh*.

S4A

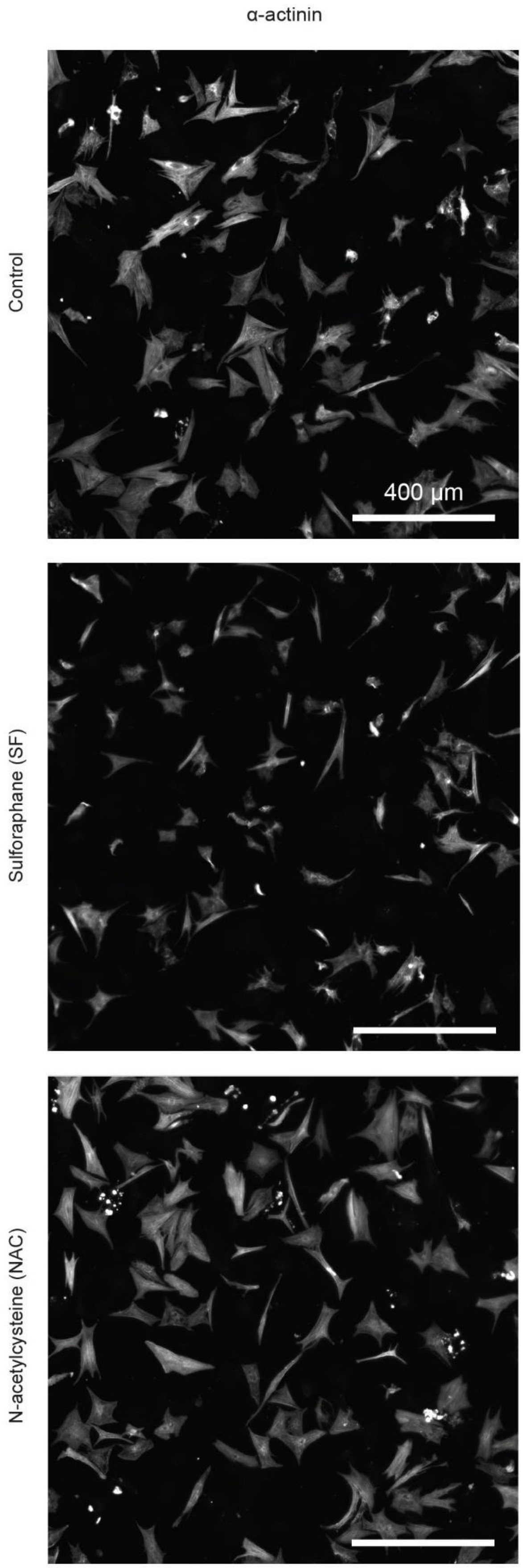
α-actinin
Control
Sulforaphane (SF)
N-acetylcysteine (NAC)
400 µm

S4B

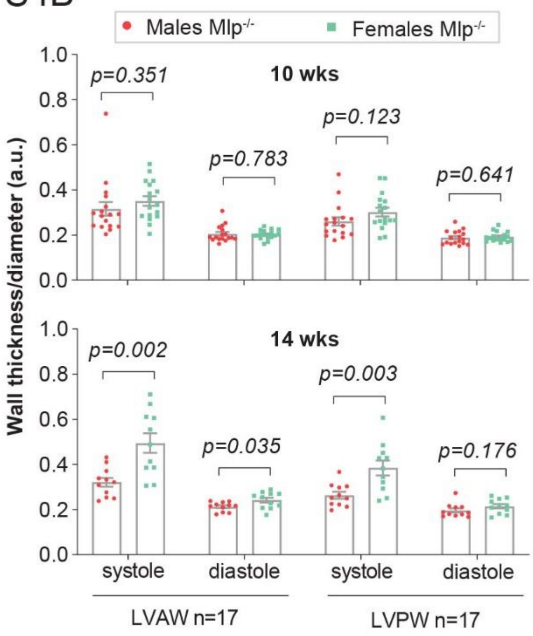
Males Mlp-/-
Females Mlp-/-
10 wks
14 wks
Wall thickness/diameter (a.u.)
p=0.351
p=0.783
p=0.123
p=0.641
p=0.002
p=0.035
p=0.003
p=0.176
systole
diastole
systole
diastole
LVAW n=17
LVPW n=17

S4C

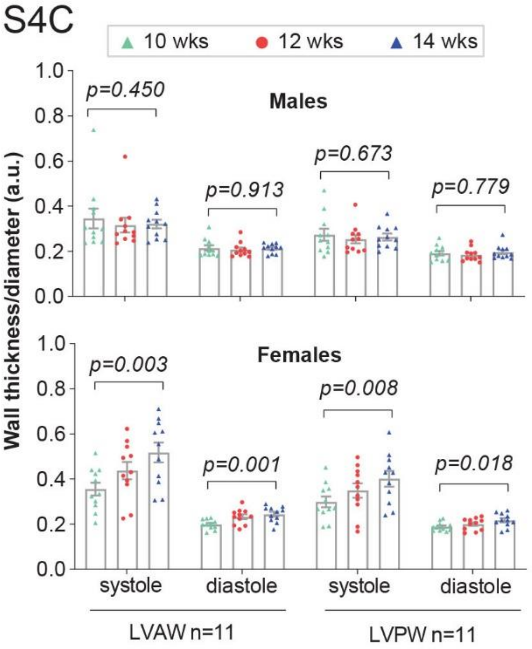
10 wks
12 wks
14 wks
Males
Females
Wall thickness/diameter (a.u.)
p=0.450
p=0.913
p=0.673
p=0.779
p=0.003
p=0.001
p=0.008
p=0.018
systole
diastole
systole
diastole
LVAW n=11
LVPW n=11

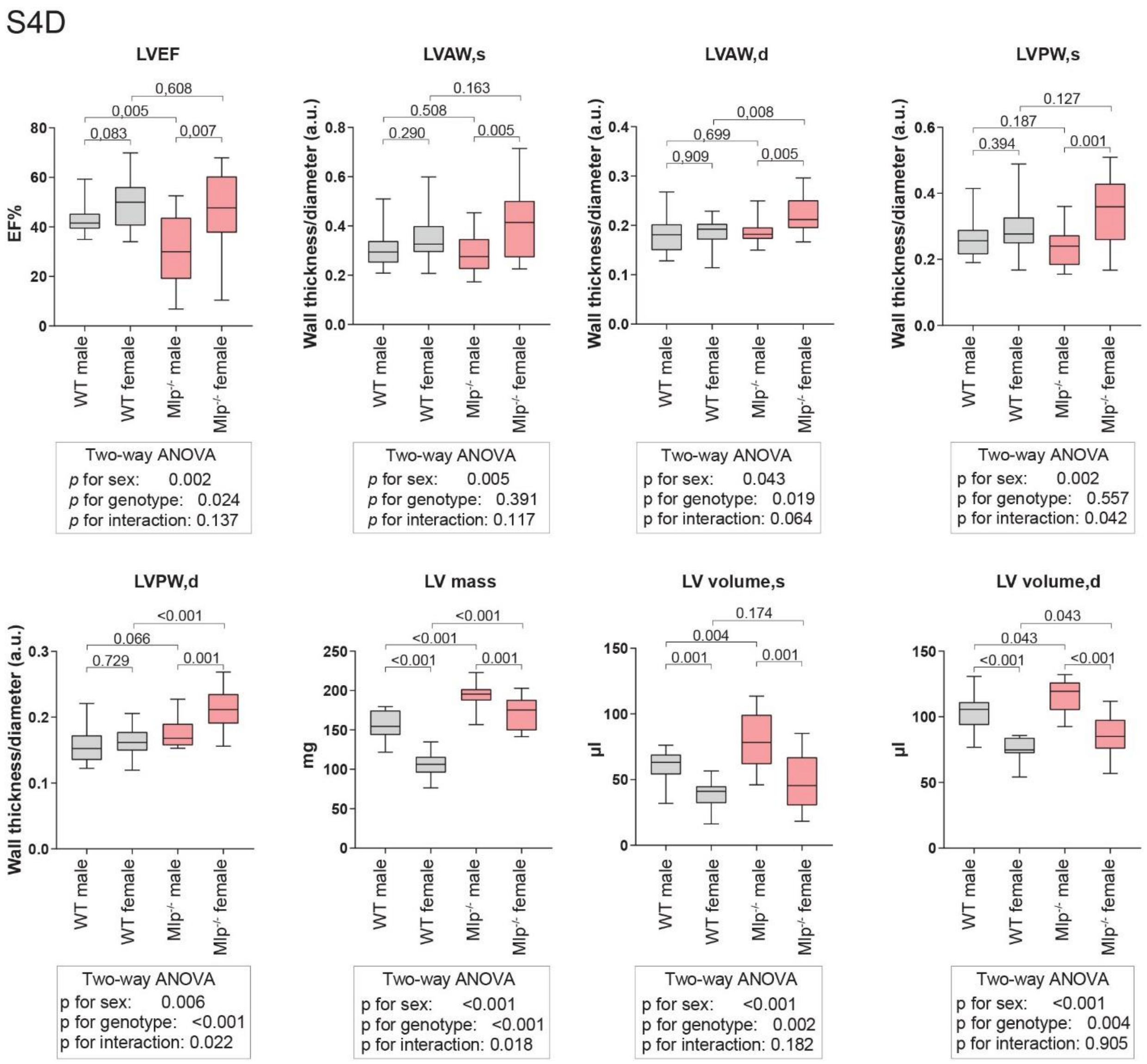

**Supplementary Figure 4: Morphology of cardiomyocytes and sex differences in responses to oxidative stress**

**54A)** The effects of NRMCs treatment with SF and NAC on cell shape: NRMCs stained for α-actinin and imaged with a bright field fluorescent microscope. These images were used for the quantifications of cell surface area and the minimum caliper diameter with ImageJ shown in figure 5E.

**54B)** LV anterior (LVAW) or posterior (LVPW) wall thickness, normalized to LV diameter, in male and female *Mlp*$^{-/-}$ mice at the age of 10 and 14 weeks. Values are the mean value ±SEM with unpaired two-tailed *t*-test. The values in the lower panel of figure S5B (14 weeks) are the same values utilized in figure 5G. It was plotted here again to be easily compared with the upper panel (10 weeks), and to visualize the differences in wall thickness between the systole and the diastole.

**S4C)** Changes over age in LV wall thickness in males and females, normalized to LV diameter. Values are the mean value ±SEM with paired *t*-test.

**S4D)** 2-way ANOVA's analysis of the effect of sex and genotype on some cardiac parameters in *Mlp*$^{-/-}$ and their littermate WT control. The displayed *p* values on the plots are unpaired two-tailed *t*-test, while the *p* values in the boxes below the plots are for the effect and interaction of the 2-way ANOVA analysis. LV(A/P)W(s/d) is left ventricle anterior (LVAW) or posterior (LVPW) wall thickness in systole (s) and diastole (d) normalized to LV diameter. The age of the animals in this figure was between 10-19 weeks old with majority of them at 12-13 weeks old.

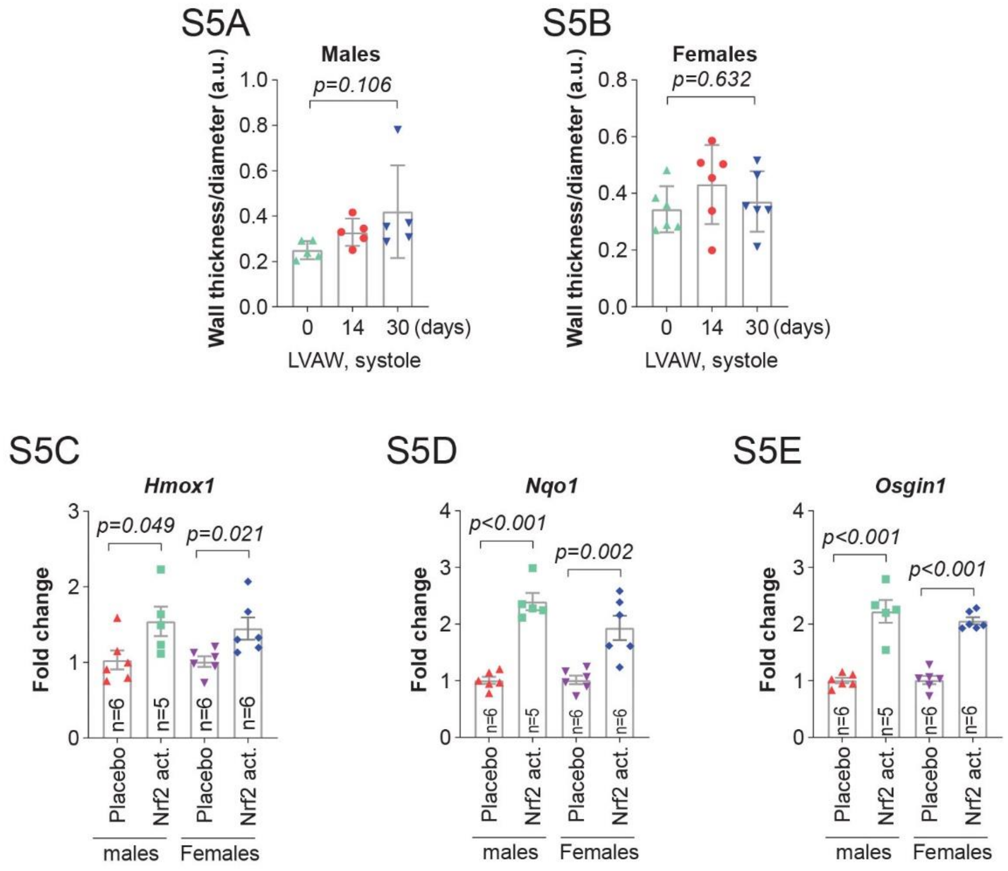

**Supplementary Figure 5: In vivo selective improvements of antioxidative treatments**

**S5A&B)** The change over time in systolic LV anterior wall (LVAW) thickness, normalized to left ventricle diameter, in male and female *Mlp*$^{-/-}$ mice during treatment with Nrf2 activator. Bars are the mean values ± SEM with paired one-tailed *t*-test.

**S5C-E)** qPCR analysis of *Hmox1*, *Nqo1*, *Osgin1* expression in the LV of *Mlp*$^{-/-}$ treated with Nrf2 activator, normalized to the vehicle treated group. Bars are the mean FC ± SEM with unpaired two-tailed *t*-test.

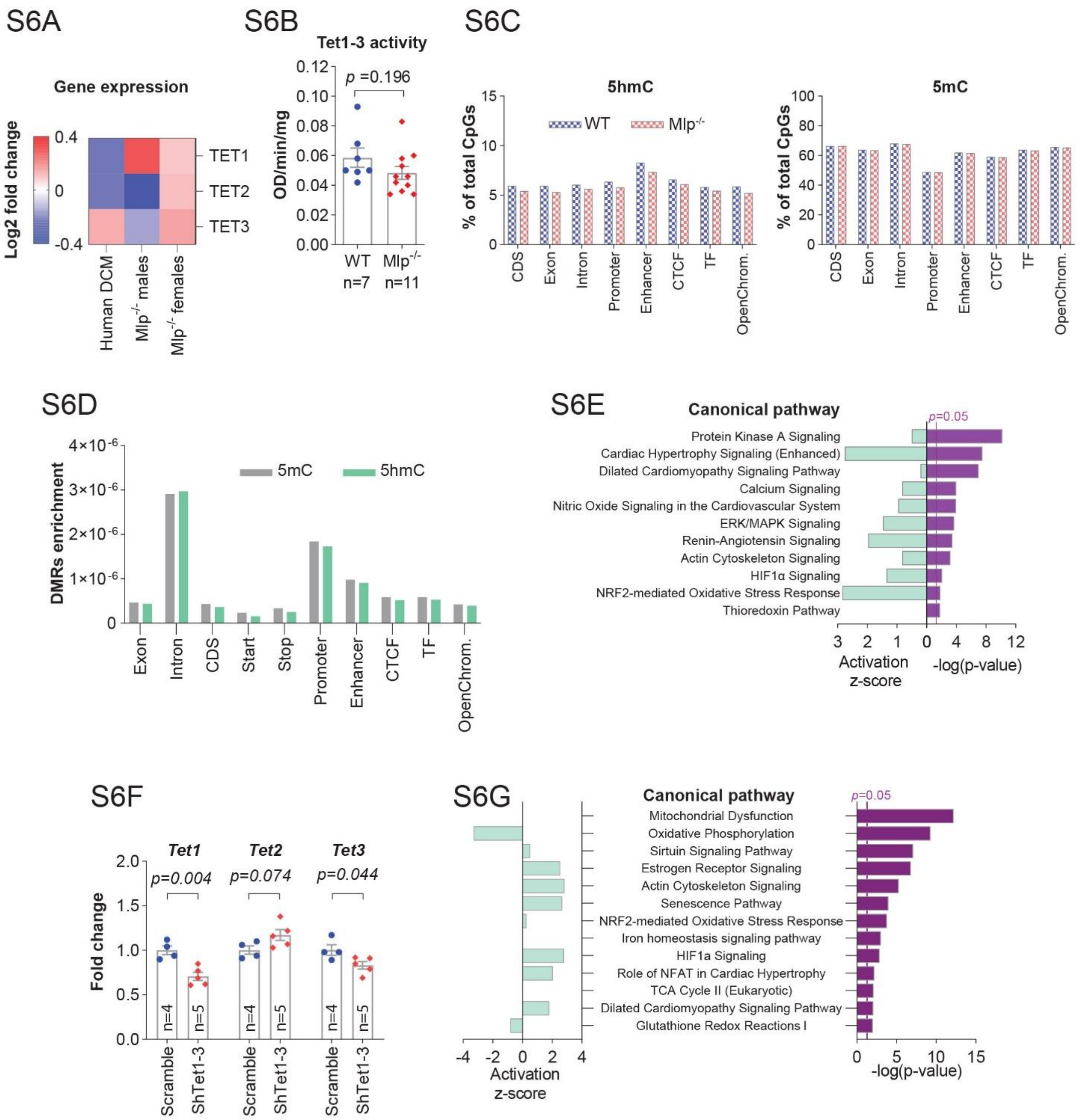

**Supplementary figure 6: Unique epigenetic control of *Idh2* expression and redox response.**

**S6A)** Expression of *TET1,2,3* genes in the LV of patients with DCM (Described in figure S1A) and male and female *Mlp*$^{-/-}$ mice analyzed by RNA sequencing.

**S6B)** The activity of Tet1-3 enzyme in the LV of male *Mlp*$^{-/-}$ mice, at the age of 12 weeks. Values are presented as the mean ±SEM with unpaired two-tailed *t*-test.

**S6C)** Percentages of 5mC or 5hmC-methylated CpGs within each functional genetic region in the myocardium of *Mlp*$^{-/-}$ mice, n=3. (CDS: Coding regions, CTCF: Transcriptional repressor CTCF binding sites, TF: transcription factors binding sites.).

**S6D)** Intersection over Union metric to visualize the enrichment of DMRs with respect to their regulatory features, in the LV of *Mlp*$^{-/-}$ mice, n=3.

**S6E)** Selected pathways from IPA on genes that harbored DMRs for 5hmC located in introns, in the LV of *Mlp*$^{-/-}$ mice

**S6F)** qPCR analysis of *Tet1-3* expression in NRCMs transduced with ShTet1-3. Values are the average of n individual wells ±SEM with unpaired two-tailed *t*-test, whereby n number is indicated on the figures.

**S4G)** Selected pathways from IPA enrichment analysis on transcriptomic data from NRCMs transduced with ShTet1-3.

Values in **S6B** and **S6F** are presented as the mean ±SEM with unpaired two-tailed *t*-test.

S7A

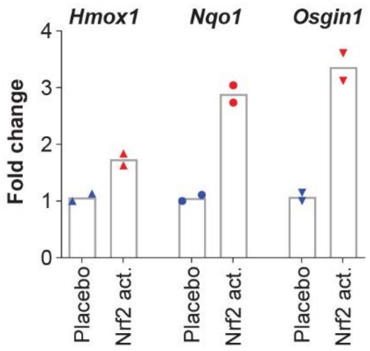

**Supplementary Figure 7: Pilot *in vivo* study with Keap1 inhibitor (AZ925)**

**S7A)** qPCR analysis of *Hmox1*, *Nqo1*, *Osgin1* levels in cardiac tissues after 9 hours post-administration of single dose of 10 mg/kg of AZ925 by oral gavage. This was a pilot experiment in accordance with the PREPARE guidelines (Smith et al., 2018), that proceeded the actual *in vivo* experiment presented in figure 5. The aim of this experiment was only to verify whether KEAP1 inhibitor would induce the expression of Nrf2-downstream target gens 9 hours post-administration of single dose of 10 mg/kg in the heart, similar to what had been observed in another organ during the process of developing this compound.

#### Extended Experimental Procedures for

### Epigenetic modulators link mitochondrial redox homeostasis to cardiac function

ElBeck et al., 2022

#### 1-Human samples

Samples from explanted human hearts with familial DCM and the matched controls were obtained from Sydney Heart Bank. The donors of the control hearts had normal ECG and ventricular function with no previous history of cardiac diseases. All samples were frozen in liquid nitrogen upon collection. The functional characteristics of these samples are provided in the supplementary table (S1) and most of the samples were utilized in multiple previous studies (Table S1). The use of human samples was approved by the local ethics committee in Stockholm (2015/559-31/2) and Human Research Ethics Committees at the University of Sydney (2016/7326), and St Vincent's Hospital (H03/118). All other data from patients utilized in this work and their corresponding characteristics were publicly available (Sielemann et al., 2020; van Heesch et al., 2019). The source of the data was cited wherever this data was used.

**Table S1: Characteristic of human samples utilized in figure 1C, and provided by Sydney Heart Bank**

| Lane in figure 1C | SHB ID | sex | Age | Mutated Gene | Clinical notes | References |
|---|---|---|---|---|---|---|
| 1 | 2.007 | Male | 22 | TTN pW14974*; | LVEF 20%, with normal coronaries | (Bos et al., 2020; Vikhorev et al., 2017) |
| 3 | 2.029 | Female | 22 | pY18923* | LVEF 13%, Normal coronaries, no myocardial infarcts, diabetic, dual chamber pacemaker. Possible viral cause. | (Vikhorev et al., 2017) |
| 5 | 2.066 | Male | 43 | OBSCN pE963K, DSP pR1537C | LVEF 10%, no ischemic heart disease. Possible viral cause. | (Bos et al., 2020; Vikhorev et al., 2017) |
| 7 | 3.133 | Female | 60 | TTN pN16499K fs*5 | LVEF 25%; Diagnosis with IDCM 5 years before transplantation. Diffuse interstitial fibrosis | (Vikhorev et al., 2017) |
| 9 | 4.100 | Male | 22 | TTN pR23464T fs*41 | LVEF 15%, no coronary artery disease. Diagnosis with possible post-viral CM, possible familial CM. | (Vikhorev et al., 2017) |
| 11 | 4.125 | Male | 37 | TTN pE23464T fs*41 | LVEF 15% Diagnosis with severe familial DCM. IHD; pleural effusion. Moderate CAD (50% in LAD) | (Vikhorev et al., 2017) |
| 2 | 7.040ies | Male | 37 | Control | Massive posterior basilar artery infarct; LVEF >55%, no significant valvular pathologies. History of epilepsy | (Brayson et al., 2019; Land et al., 2017; Martin-Garrido et al., 2018) |
| 4 | 5.138 | Male | 23 | Control | Hypoxic brain injury 2° to self-strangulation Subarachnoid haemorrhage; clean coronary arteries. Had a cardiac standstill for 13 minutes, then restarted. No coronary disease | (McNamara et al., 2017; Mollova et al., 2013; Vikhorev et al., 2017) |
| 6 | 4.131 | Male | 45 | Control | Hypoxic brain injury | |

| | | | | | | |
|---|---|---|---|---|---|---|
| 8 | 8.010 | Female | 43 | Control | CoD L middle cerebral artery bleed and aneurism. Parenchymal haematoma. Heart was perfused with formalin via coronary aa. | (McNamara et al., 2017) |
| 10 | 5.090 | Female | 42 | Control | Subarachnoid haemorrhage while driving, Middle cerebral artery bleed-clean coronary artery, heart still beating at harvest | |
| 12 | 6.052 | Male | 48 | Control | Spontaneous intracranial haemorrhage, No cardiac arrest | (Mollova et al., 2013) |

**2-Animal experiments**

Animal experiments were performed in accordance with European ethical regulation (Directive 2010/63/EU) and approved by local animal ethics committee of Linköping (permit numbers S43-15, 1369 and 2713) in Sweden, and the responsible government agency of Unterfranken (RUF-55.2.2-2532-2-659) in Germany. Animals were housed on a 12 h light/12 h dark cycle with free access to chow and water.

**2-1-Animals**

The background of muscle lim protein-deficient mice ($Mlp^{-/-}$), also known as cysteine and glycine rich protein 3 ($Csrp3^{-/-}$) is a hybrid cross of original 129/Sv background (Arber et al., 1997) with C57BL/6N strain. The mice were bred inhouse for multiple generations. Heterozygous males and females were bred to obtain both WT and $Mlp^{-/-}$ animals. Both male and females were utilized for experiments at the age of 10-14 weeks. If animals of other ages were used, detailed descriptions were mentioned in the figure legends.

Rats from Sprague Dawley strain were purchased from Charles River Laboratories and bred inhouse to obtain neonatal pups.

**2-2-In vivo KEAP1 inhibitor study**

Novel Keap1 inhibitor (AZ925) and the vehicle control were formulated and provided by AstraZeneca- MedImmune. The study was designed and performed in accordance with the PREPARE guidelines (Smith et al., 2018). The dose utilized (10 mg/kg) was adapted from previous unpublished studies performed during the development of AZ925. We validated this dose through a pilot study, by administrating 10 mg/kg of AZ925 by a single oral gavage in two mice. After 9 hours post-administration, we terminated the study and assessed the induction of Nrf2- targeted genes (Hmox1, Nqo1 and Osgin1) by qPCR analysis in the heart, liver, brain, and kidneys. We observed a robust induction of these genes (1.5-7-fold increase) in all examined organs. The data of the pilot study from heart is shown in figure S7.

*Mlp*$^{-/-}$ animals at the age of 10 weeks were treated by a daily gavage with 10 mg/kg of AZ925 or vehicle for 30 days. Both male and female mice were included in the study, with 6 animals in each of the 4 groups (Keap1 inhibitor/Placebo-Males/females). The mice were randomized to different treatment groups based on their body weight. Echocardiography was performed at days 0, 14 and 30 (see Figure 5A for experimental scheme). One male in the Keap1 inhibitor group died immediately after first gavage, but all other animals survived to the end of the study, with no observable changes in their behaviors or body weight. The investigators were blinded to the different groups during experimental procedures.

**2-3-Echocardiography**

Echocardiography was performed with the ultrasound Vevo 3100 System, with the MX550D probe (Visualsonics, Canada). Animals were initially anaesthetized with 3% isoflurane (Forene®) mixed to oxygen (Oc-P50, Sysmed) and then maintained at 1.5% isoflurane during the examination. The animals were kept on a Physio Plate (Visualsonics, Canada) during whole procedure to maintain normal body temperature, and to monitor ECG and respiration. Body temperature was monitored by an inserted rectal probe. An external heat lamp was used when it was needed. Left ventrical parasternal long and short axis projections were captured in both B-Mode and motion-mode (M-MODE). Short axis midventricular level, with a clear projection of the papillary muscles, was captured after rotating the probe ~90° clockwise from the long-axis view. By using the tool *LV Analysis* in the Vevolab software (version 5.5.0), several cardiac parameters were evaluated from the M-mode's short-axis view, in at least three beats and averaged. Among the main obtained parameters were left ventricular internal diameter (LVID) in systole (s) and diastole (d), LV's ejection fraction, and the left ventricular posterior and anterior wall thickness (LVPW, LVAW) in systole and diastole. The thickness of anterior or posterior LV wall was normalized by dividing the inferred wall thickness in diastole or systole to the measured LV diameter in diastole or systole, respectively. The investigators were blinded to the groups during experimental procedures.

**2-4-Tissue collection**

Mice hearts were excised immediately after cervical dislocation, washed with cold PBS to remove the blood and then weighed. Left ventricles were quickly separated and flash frozen in liquid nitrogen. Mice included in the Keap1 inhibitor study were anaesthetized (10 ± 3 hours post-administration of the last dose) with 3% isoflurane (Forene®) and then the hearts were excised and processed as described.

**2-5-Tissue grinding and homogenization**

Frozen human and murine left ventricle cardiac tissues were grinded and homogenized in Cellcrusher (Cellcrusher, Irland) in liquid nitrogen.

**3-Experiment on isolated Neonatal rat cardiomyocytes (NRCMs)**

**3-1 NRCMs isolation**

Hearts from 3 days old Sprague-Dawley rats were excised after decapitation and placed immediately in ice-cold preservative buffer (concentrations in mM: NaCl 13, KCl 0.54, HEPES 2.5, $MgCl_2$ 0.05, $NaH_2PO_4$ 0.04, Glucose 22.20, pH was adjusted to 7.4 with NaOH). Ventricles were separated from atria and minced into small pieces. Tissue pieces were dissociated into single cells at 37$^oC$ by utilizing the gentleMACS Octo Dissociator with Heaters (Miltenyi Biotec, Bergisch Gladbach, Germany) using Neonatal Heart Dissociation Kit (Miltenyi Biotec) according to the manufacturer's instructions. The single cell suspension was diluted with 7.5 ml of the cardiomyocytes plating media (CM plating media: DMEM (high glucose): M199 (4:1), supplemented with 5% horse serum, 2.5% fetal bovine serum, 20 mM HEPES and 100 U/ml penicillin/streptomycin (Thermo Fisher Scientific, Waltham, Massachusetts, USA)) at room temperature (RT) to inactivate the digestive enzyme mix and to gradually bring the temperature of the cell suspension down, and then passed through MACS Smart Strainers (70 µM, Miltenyi Biotec) to remove tissue residues. Then the red blood cells were lysed with Red Blood Cell Lysis Solution (Miltenyi Biotec). The lysis step was performed at RT, but subsequent enzymatic treatment was done in ice-cold PBS buffer. After the red blood cell lysis, cardiomyocytes were enriched in the cellular suspension by allowing non-cardiomyocytes to adhere to a surface of 10 cm plates (Corning) by pre-plating them for 75 min at 37$^oC$ in CM plating media. Enriched cardiomyocytes were counted with TC20 automated cell counter (BioRad, US) and plated in either 6-well or 24-well plates (Corning) according to the experiments described below. 1.25 $x10^6$ or 3.33 $x10^5$ live cells were plated in each well of the 6-well or 24-well plates, respectively. Before plating the cells, the plates were gelatinized with 0.15% gelatin solution for 1h at 37$^oC$. Plated cells were maintained in incubators at 37$^oC$ with 5% $CO_2$.

**3-2 Treating NRCMs with redox active compounds**

Enriched NRCMs were plated in CM plating media in either 6-well plate for microscopic florescence imaging, or in 24-well plate for RNA extraction. After 24 h of incubation, the cells were washed once with PBS to removed dead cells and incubated for another 48 h with CM maintaining media (similar composition of the CM plating media but supplemented with 2.5 µM of the mitotic inhibitor cytarabine, (Sigma-Aldrich, Germany)). After the 48 h incubation in the maintaining media, cells were treated with hydrogen peroxide, Sulforaphane or N-Acetyl-L-cysteine (Sigma-Aldrich, Germany).

**3-2-1-Hydrogen peroxide ($H_2O_2$)**

A stock solution (50 mM) was prepared in water and diluted with NRCM maintaining media to concentrations of 25, 50, 100, 200 or 300µM. NRCMs were treated with $H_2O_2$ for 24 h, thereafter, the media was removed and exchanged with a fresh media without $H_2O_2$ for another 24 h. Cells were collected at different time-points at 6, 24 or 48 h from the start of the treatment (see Figure.2D for the experimental scheme).

**3-2-2 DL-Sulforaphane (SF) and N-Acetyl-L-cysteine (NAC)**

SF powder was dissolved in DMSO to obtain a 0.1 mg/µl solution, which was aliquoted and stored at -20$^{o}$ C. This solution was diluted with water to get 500 µM stock solution, which was further diluted with NRCMs maintaining media to a final concentration of 5 µM. SF solution were freshly prepared for every experiment from the frozen stock.

Fresh stock solution of 100 mM NAC was prepared from NAC powder and diluted with NRCM maintaining media to a final concentration of 3 mM.

NRCMs were treated with SF or NAC for 48 h or 72 h. The maintaining media was changed every 24 h with fresh supplements of SF or NAC. For RNA expression analysis, cells were collected after 6, 24 or 48 h of treatment (see Figure 4A for the experimental scheme). For immunofluorescence cells were treated for 72 h and fixed with 4% formalin for 10 min at 37$^{o}$C, then washed twice with PBS (See section 5-1).

**3-3- Transducing NRCMs with ShRNA targeting L2hgdh, D2hgdh or Tet1-3**

All constructs were AAV based vectors packaged into AVV9 viruses (VectorBuilder) where the small RNA expression was driven by the Pol III promoter. Enhanced green fluorescent protein (EGFP) transcript was added to the construct under the cytomegalovirus (CMV) promoter.

Two experimentally validated ShRNA sequences targeting each of the rat's *Tet1-3* were obtained from published literatures (see Table S2). To achieve simultaneous knockdown of all *Tet1-3* in individual cardiomyocytes, three ShRNA sequences, each targeting one of the three *Tet1-3* mRNA were constructed together in one vector, resulting in two vectors. For targeting rat's *L2hgdh* and *D2hgdh* mRNA, we tested multiple predicted ShRNA sequences. Each designed vector contained two or three sequences targeting either *L2hgdh* or *D2hgdh*, to increase the efficiency of finding a positive hit (Table S2). The vector that achieved more than 50% knockdown of the target's mRNA at a dose of 10000 multiplicity of infection (MOI) was selected for downstream experiments and their ShRNA sequences are shown in table S2., and their full map are shown in the supplementary file F1. Enriched cardiomyocytes were plated in CM plating media in either 6-well plate for protein extraction, or in 24-well plate for RNA extraction. After 24 h of plating, the cells were washed twice with PBS, and transduced with AAV9 in CM maintaining media. To increase the transduction efficiency, cells were kept in the same media for 3 days. After 3 days, one volume of the fresh maintaining media was added to each well. Then, at days 4 and 5 the media was exchanged with a fresh maintaining media. Cells were collected after 6 days of transduction.

**Table S2: Sequenced of ShRNA utilized in the current work**

| Construct ID | Target gene | ShRNA sequences | Map | References |
|---|---|---|---|---|
| **1038dup** | Tet1-3 | Tet1: GGAGGGATTTCTCACGTTA<br>Tert2 GGATGTAAGTTTGCCAGAAGC<br>Tet3: GCTCCAACGAGAAGCTATTTG | https://en.vectorbuilder.com/vector/VB191127-1038dup.html | (Zhao et al., 2014) |
| **1073xvx** | Tet1-3 | Tet1: GCTCATGGAGACTAGGTATGG<br>Tet2: CTCAGGGATGTCCTATTGCTAAA<br>Tet3: AAGCGCAACCTATTCTTGGAA | https://en.vectorbuilder.com/vector/VB191127-1073xvx.html | (Hsieh et al., 2016; Zhao et al., 2014) |
| **1186ueu** | L2hgdh | AGTAAGGATGGGATGAAATAT,<br>GGAGTCACTGAAAGCTAAATT | https://en.vectorbuilder.com/vector/VB200603-1186ueu.html | - |
| **1008zgf** | D2hgdh | ACGTGTTCAAGTATGACTTAT,<br>TTGGTGCCTTGGAGCCTTATG,<br>CTGAACTGCCTGACCTCTTTG | https://en.vectorbuilder.com/vector/VB200529-1008zgf.html | - |
| **1020znr** | Scramble | CCTAAGGTTAAGTCGCCCTCG | https://www.vectorbuilder.kr/vector/VB180117-1020znr.html | - |

#### 4-Metabolites analyses

L-malate, succinate, lactate and L/D2HG were analyzed by a targeted mass spectrometric approach. We developed an analytical method that enables extraction and separation of the two 2HG enantiomers in heart tissues and NRCM cells, based on a previously published method for analyzing L/D2HG in cancer tissues (Cheng et al., 2015). The method utilizes chemical derivatization of L/D2HG by N-(p-toluenesulfonyl)-L-phenylalanyl chloride (TSPC). TSPC was reported to enable higher sensitivity for L/D2HG detection and quantification than DATAN based method (Cheng et al., 2015).

##### 4-1-Metabolites extraction

Heart tissues: 10 mg of homogenized and grinded tissues were weighed in a 2 ml Eppendorf tube (Sarstedt). Then 100 µl of deionized water containing 0.5 nmol of heavy isotope-labelled L/D2HG internal standard (IS) (Deuterated RS2HG, Cambridge Isotope Laboratories) was added to each tube, with a 0.5 mm metallic ball (Qiagen). The tubes were immediately homogenized by the TissueLyser LT (Qiagen, Germany) at 4°C for 2 min at 50 Hz and then spun quickly before placing them back on dry ice. After the tubes' content got completely frozen, the tubes were allowed to thaw partially, and then homogenized again for 2 min at 50 Hz. This cycle of freezing, thawing, homogenization, and brief spinning was repeated for the total of 4 times to break down all cellular and mitochondrial walls. The temperature was always kept below 4°C during the whole procedure to minimize any enzymatic reaction of the metabolites. After last homogenization cycle, 400 µl of cold pure methanol (-80°C) were added to each tube (to obtain 80% MeOH aqueous solution) to extract the soluble metabolites, and the tubes were homogenized again for another 2 min at 50 Hz, and then incubated on dry ice for 5 min. As negative controls, 3 tubes containing 100 µl of deionized water with 0.5 nmol of deuterated-RS2HG, L2HG or D2HG standards underwent the same described procedure with the samples.

NRCMs: Transduced NRCMs with ShRNA targeting *L2hgdh*, *D2hgdh* or scramble control were trypsinized with TripleE (Invitrogen), centrifuged at 6000 g for 10 min at 4°C to remove the supernatant, and then stored at -80°C. Then 40 µl of water containing 0.5 nmol of the IS deuterated-RS2HG was added to each pellet of cells. The pellets were thawed on ice and then frozen again for a total of 10 cycles, with vortexing in between, to completely lyse the cells. Then 160 µl of -80°C methanol was added. The mixture was vortexed and incubated on dry ice for 5 min.

**4-2- Derivatization with TSPC**

TSPC was dissolved in acetonitrile (ACN) for a stock concentration of 62.5 mM, and then aliquoted and stored at -80°C.

The 80% MeOH aqueous extracts from tissue and NRCMs were centrifuged at 4°C at the maximum speed of 20800 *g* for 10 min. The supernatant was transferred to a new tube and the solvent was completely evaporated under vacuum at 45°C (SpeedVac Savant SPD2010). The dried metabolites' residues were dissolved in 100 µl of acetonitrile by vigorous vortexing. Then another 60 µl of ACN containing 2 µl of pyridine was added. The derivatization reaction was initiated by adding 3.2 µl of 62.5 mM TSPC and incubating the tubes at 25°C for 10 min. The solvent was then evaporated completely under vacuum at 45°C (SpeedVac Savant SPD2010). The resulting precipitants were dissolved with 15 µl 50% ACN/water and centrifuged at a maximum speed of 20800 g for 10 min. ~14 µl were transferred to a 0.2 mL PCR strips and stored at -80°C for the mass spectrometric analysis.

**4-3-Mass spectrometric analysis**

Two µl of samples were injected in an Ultimate™ 3000 UPLC coupled with a heated electrospray ion source to an Orbitrap™ Fusion Lumos™ tribrid mass spectrometer (ThermoFisher Scientific). The chromatographic separation was achieved using a 25 cm long (2.1 mm i.d., 5 µm particle size) Inertsil™ ODS-3 column (GL Sciences, Tokyo, Japan) at 35°C. Formic acid in water (0.1%, as solvent A) and a 50:50 mixture of acetonitrile and methanol (as solvent B) were employed as the mobile phase. A gradient of 3 min 30% B, 7 min 30–70% B, 15 min 70% B, 1 min 70–30% B, and 14 min 30% B was used at a flow rate of 200 µl/min.

The mass spectrometer was set to acquire tandem mass spectra in small molecule mode with negative ion detection. The precursors defined in an inclusion mass list were quadrupole isolated aiming at minimum six points across the peak with *m/z* 0.7 isolation width in the ranging from *m/z* 50 to 500 at a resolution of R=30,000 (at *m/z* 200) targeting $5x10^4$ ions for maximum injection time of 54 ms, using higher energy collision dissociation (HCD) fragmentations at 27% normalized collision energy in 2 s cycle time.

Skyline v20.1.0.155 (MacCoss Lab, Dept. of Genome Sciences, University of Washington (Adams et al., 2020)) was used for data analysis by importing raw data files with target masses, including 2HG labelled with heavy isotope ($\Delta_{mass}$=3 Da). The ion match tolerance was set to 0.05 *m/z*, allowing automatic selection of all matching transitions. The imported data was manually controlled for peak boundaries. The peak areas were extracted and used for quantitative comparisons. When multiple fragments were detected from the precursor ions, the fragment with the highest intensity were utilized for the quantitative comparisons (Table S3).

**Table (S3): Inclusion mass list and fragments mass**

| Compound Name | Formula | Adduct Positive | *m/z* | z | t (min) | | HCD Collision Energy (%) | Fragment 1 | | Fragment 2 | |
|---|---|---|---|---|---|---|---|---|---|---|---|
| | | | | | start | stop | | *m/z* | Formula | *m/z* | Formula |
| **Lactic acid** | $C_3H_6O_3$ | -H | 89.0244 | 1 | 0 | 30 | 27 | 71.0139 | $C_3H_3O_2$ | | |
| **SA** | $C_4H_6O_4$ | -H | 117.0193 | 1 | 2.53 | 4.53 | 27 | 73.0295 | $C_3H_5O_2$ | 99.0088 | $C_4H_3O_3$ |
| **2HG-TSPC** | $C_{21}H_{23}NO_8S$ | -H | 448.1072 | 1 | 16.23 | 19.23 | 27 | 318.0806 | $C_{16}H_{16}NO_4S$ | 155.0172 | $C_7H_7O_2S$ |
| **2HG-TSPC_heavy** | $C_{21}H_{20}D_3NO_8S$ | -H | 451.126 | 1 | 16.23 | 19.23 | 27 | 318.0806 | $C_{16}H_{16}NO_4S$ | 155.0172 | $C_7H_7O_2S$ |
| **MA-TSPC** | $C_{20}H_{21}NO_8S$ | -H | 434.0915 | 1 | 19.16 | 21.16 | 27 | 318.0806 | $C_{16}H_{16}NO_4S$ | 155.0172 | $C_7H_7O_2S$ |

#### 4-3-Data analysis and calculations

The levels of L/D2HG in the LV myocardium of WT and *Mlp*$^{-/-}$ mice were compared by dividing the peak areas of the 155 *m/z* fragments of L/D 2HG-TSPC to the corresponding peak areas of the spiked-in L/D2HG-TSPC_heavy. The resulting values were then normalized to tissue weight. The levels of L/D2HG in transduced NRCMs with ShL2hgdh and ShD2hgdh were compared to the scramble control-transduced cells by dividing the peak areas of the 318 *m/z* fragments of L/D 2HG-TSPC to the corresponding peak areas of the spiked-in L/D2HG-TSPC_heavy. The levels of L-malate (MA), succinate (SA), and lactate in cardiac tissues of WT and *Mlp*$^{-/-}$ mice were compared by normalizing the peak areas of the 155 *m/z* fragment of MA-TSPC, 73 *m/z* fragment of succinate and 71 *m/z* fragment of MA-TSPC, respectively to tissue weight.

**4-4 2-oxoglutarate (2OG) quantification**

The levels of 2OG in the LV myocardium of WT and $Mlp^{-/-}$ male mice were measured by utilizing the colorimetric 2-oxoglutarate Assay Kit MAK054 (Sigma-Aldrich) according to the manufacturer's instructions with some modifications. In brief, 20 mg of homogenized and grinded frozen heart tissues were weighed in a 2 ml Eppendorf tube (Sarstedt). Then 100 µl of the assay buffer was added to each tube, with a 0.5 mm metallic ball (Qiagen). The tubes were immediately homogenized by the TissueLyser LT (Qiagen, Germany) at 4$^{o}$C for 2 min at 50Hz and then spun quickly before placing them back on dry ice. After the tubes' content got completely frozen, the tubes were allowed to thaw partially, and then homogenized again for 2 min at 50Hz. This cycle of freezing, thawing, homogenization, and brief spinning was repeated for the total of 4 times to break down all cellular and mitochondrial walls. The temperature was always kept below 4$^{o}$C during the whole procedure to minimize any enzymatic reaction of the metabolites. Then the tubes were centrifuged at 4$^{o}$C at the maximum speed of 20800 g for 10 min, and then the supernatant was transferred to a new tube and centrifugation was repeated for another 15 min. 50 µl of the supernatant was transferred to a 0.2 mL PCR strips and stored at -80$^{o}$C. Two µl of the tissue's extract was diluted with NP40 buffer and utilized for estimating total protein concentration by BCA assay. To measure 2OG quantity, we utilized a standard curve with the concentrations of 0, 0.5, 1, 1.5, 2, 2.5 nmol/well. Samples and standards were measured in duplicates, in a total volume of 25 µl per well of 384-well plate (Corning). The absorption was measured at 570nm in SpectraMax® i3 (Molecular Devices, US). The measured quantity of 2OG was normalized to the total protein content and plotted.

**5-Imaging**

**5-1-Immunostaining**

NRCMs cultured in 6-well plate (Corning) with a plastic bottom were fixed with 4% buffered formalin as described in section (3-2-2). Cells were washed with PBS and then permeabilized with 0.2% Triton X-100 for 10 min at RT and blocked with a blocking buffer (1% BSA, 22.52 mg/ml glycine and 0.1% Tween 20 in PBS) for 30 min at RT. The primary antibody (mouse anti-α-actinin antibodies A7811, Sigma-Aldrich) were diluted with an incubation buffer (1%BSA and 0.1% Tween 20 in PBS,1:1000) and incubated overnight at 4$^{o}$C. Secondary antibodies (goat anti-mouse, Alexa Fluor 568-conjugated antibody ab175473, Abcam) was diluted with incubation buffer (1:1000) and incubated for 1h at RT. Cells were washed 3 times with PBS after each step. Confocal and widefield images were acquired as it described in the following section.

**5-1-1-Microscopy imaging**

Images were acquired with a S Plan Fluor Ph2 ELWD 60x/0.70 objective on a Nikon Ti2 microscope equipped with a CREST Optics V3 spinning disk confocal (50 um pinholes). The emission iris was closed to match the objective, according to the specifications by CREST Optics. The ring of the objective was adjusted to the thickness of the plate bottom. The camera used was a Photometrix BSI Express Back illuminated sCMOS (for widefield images; Sensitivity 11 Bits mode, readout speed 200 MHz) or a Photometrix 95B Back illuminated sCMOS (for confocal images. Sensitivity 12 Bits mode).

The α-actinin channel was acquired using a 546 nm laser and a Pentaband emission filter (441/30; 511/26; 593/37; 684/34; 817/66 nm). The laser power and exposure time were set to get the brightest possible images without any saturation. The same settings were used for all the samples to be compared.

The brightness and contrast of the analyzed images were adjusted to the same settings, and the images were acquired either in wide field as tiles (9x9) or with the spinning disk confocal. Only the wide field images were further analyzed.

**5-1-2-Image analysis**

Image analysis was performed with the help of the SciLifeLab BioImage Informatics Facility. In short, we developed an unbiased approach to profile the shape of neonatal rat cardiomyocytes in Fiji (Schindelin et al., 2012). The α-actinin channel was utilized for segmentation. The workflow consisted of a pre-processing step in which the median filter was applied to the input images in order to reduce noise. Then, the images were segmented using the Li's Minimum Cross Entropy thresholding method (Li and Lee, 1993; Li and Tam, 1998; Sankur, 2004). To avoid any bias in quantifying segmented cardiomyocytes between different treatments, we excluded all cropped cells that were on the images' edges. We also excluded segmented particles with areas smaller than 50000 pixels.

To quantify the changes in cell shape upon treatments, we utilized the minimum caliper diameter (MinFeret shape descriptor in Fiji), which measures the smallest diameter between any two points at the boundary of a segmented object (Schindelin et al., 2012) . We also utilized the area of the segmented objects to visualize changes in cardiomyocytes' size upon different treatments. However, NRCMs in 2D culture acquires variety of random shapes and sizes, and the diversity in the shapes and sizes were further greatly enhanced by the SF and NAC treatments, which also affected the identifications of contiguous cells' boundaries. This fact made it impossible to segment all individual cardiomyocytes, especially in the NAC treatment, where cells were hypertrophied and more intertwined. This problem was not even possible to resolve by manual curation of cells' borders or by staining the cytoplasmic membrane (data not shown). As our interest was to quantify the changes in cell area and diameter between different treatments, we segmented connected cells as one object. We segmented 6 images from the control group, and 8 images from each of SF and NAC treatments. The segmentation resulted in 123-196 objects per image. For each object of each image, we computed the area and minimum diameter. To plot them, we pooled the measures of each condition together. Pooling them resulted in groups of values, in which, each value originated from one image (6 for the control condition,8 for NAC and 8 for SF treatments). Then we calculated the average of each group. This resulted in 148 objects in the control condition, 144 objects in the NAC treatment, and 196 objects in the SF treatment. These averaged values of the segmented objects' area and MinFeret were plotted in GraphPad Prism. The implemented image analysis pipeline is available in Github. All quantified raw images will be available on Image Data Resource (IDR) (https://idr.openmicroscopy.org/).

##### 5-2 Transmission electron microscopy (TEM)

Small pieces (2-3 $mm^3$) of LV myocardium of WT and *Mlp*$^{-/-}$ mice were dissected and fixed for 30 minutes at room temperature in fixation buffer (2% glutaraldehyde + 1% paraformaldehyde in 0.1 M phosphate buffer, pH 7.4) and stored at 4$^{o}$C. After rinsing them with 0.1 M phosphate buffer, pH 7.4, specimens were postfixed for 2 h in 2% osmium tetroxide 0.1 M phosphate buffer, pH 7.4 at 4$^{o}$C, then dehydrated in ethanol followed by acetone and embedded in LX-112 (Ladd, Burlington, Vermont, USA). Leica ultracut UCT (Leica, Wien, Austria) was utilized to prepare ultrathin sections (approximately 50-60 nm). The sections were later contrasted with uranyl acetate followed by lead citrate and examined in a 100 kV Hitachi HT 7700 (Tokyo, Japan) and the digital images were captured with a Veleta camera (Olympus Soft Imaging Solutions, GmbH, Münster, Germany). Mitochondrial volume density (Vv) was calculated by point counting on printed digital images using a 2 cm square lattice. From each animal, 9 randomly taken images were utilized for appropriate sampling. The number was estimated by utilizing the cumulative mean plot (Weibel et al., 1979).

**6-Biochemical assays:**

**6-1-Mitochondrial respiration:**

Mitochondria were isolated from fresh WT and *Mlp*$^{-/-}$ adult hearts (12-15 weeks old). Oxygen consumption of isolated mitochondria was measured with the exact same experimental procedures and settings described in a previous work (Nickel et al., 2015).

**6-2- Determining enzymatic activity of Idh2:**

Idh2 activity were measured from mitochondria isolated from frozen LV myocardium of WT and *Mlp*$^{-/-}$ mice (males, 12 weeks old). The isolation procedure for mitochondria was adapted from (Nickel et al., 2015) with some modifications. Grinded frozen cardiac tissues (5mg) were weighed in a 2 ml Eppendorf tube (Sarstedt) and homogenized with 200 µl of the isotonic isolation buffer (IS; in mM: sucrose 75, mannitol 225, HEPES 2, EGTA 1, pH 7.4, 4$^{o}$C). The homogenization was performed in the TissueLyser LT (Qiagen, Germany) for 2x2 min at 50Hz with the presence of one 0.5 mm metallic ball (Qiagen) and two small 0.2 mm metallic beads (Retsch). After a brief spinning, additional 400 µl of the isolation buffer was added and homogenized again for 2x2 min at 50Hz. The homogenate was centrifuged at 480 g at 4$^{o}$C for 5 min, and a 400 µL of the upper supernatant was transferred to a new tube which was kept on ice. The homogenization step (4 min at 50Hz) was repeated with a fresh 400 µl of isolation buffer, and then 400 µl of the supernatant was combined with previous fraction. The remaining cellular pellet was utilized to obtain the nuclear extract (See section 6-3). To remove any cellular debris from the supernatant containing the mitochondria, the tube was gently flicked and centrifuged again for 480g for 5min, then 700 µL of supernatant was moved to a new tube without disturbing the pellet. The supernatant was further centrifuged at 7700 g for 10 min at 4$^{o}$C to obtain mitochondrial pellet. Mitochondrial pellet was washed twice with 200 µl of isolation buffer without ETGA (mitochondrial suspension solution MMS). The pellet was lysed in 50 µl of MMS by three cycles of freezing and thawing and then stored at -80$^{o}$C. Temperature was always kept below 4$^{o}$C during the whole isolation procedure. 5 µl of mitochondrial lysate was diluted with NP40 buffer and utilized for total protein quantification by BCA assay.

The activity of Idh2 was measured with the same reaction mix described in (Nickel et al., 2015) (in mM: Tris–HCl 10 [pH 8.0], $NADP^+$ 0.2, $MgCl_2$ 5, and isocitrate 2) in 384-well plate with a total volume of 80 µl per well. The change in the absorption at 340 nm was continuously recorded for 60 min with the kinetic function of SoftMax Pro 7 in SpectraMax® i3 plate reader (Molecular Devices, US). All samples were measured in duplicates. As a negative control, Idh2 was inactivated in the samples by adding 1 mM of N-ethylmaleimide (NEM) to the reaction mix (Smyth and Colman, 1991). 5 µg of mitochondrial proteins were utilized per each reaction. The generated amounts of NADPH were estimated by using Beer–Lambert law (A=εbC), where A is the measured absorbance at 340 nm, ε is the molar attenuation coefficient (ε= 6.22 mM-1 cm-1 for NADPH), b is the length of the light path (0.7 cm) and C is the concentration of generated NADPH.

**6-3- Determining enzymatic activity of Tet1-3:**

The *ex vivo* enzymatic activities of Tet1-3 were compared between the WT and the *Mlp*$^{-/-}$ male mice (12 weeks old) by utilizing the colorimetric Epigenase 5mC-Hydroxylase TET Activity Assay Kit (P-3086, Epigentek) according to the manufacturer's instructions. The nuclear extract was prepared by utilizing the nuclear extraction kit (OP-0002, Epigentek). The input materials for nuclear protein's extraction were the remaining cellular pellets in section (**6-2**) after isolating subsarcolemmal mitochondria. 15 µg of extracted nuclear proteins were used per well for the measurement of TET activity. Samples were measured in 4 replicates.

**7- Molecular biology**

**7-1-Western blotting:**

Tissues: 10 mg of grinded LV myocardium were weighed and lysed with 200 µl of modified RIPA buffer (50mM TRIS, 150mM NaCl, 1% sodium deoxycholate, 1% Sodium dodecyl sulfate, 1% Tritonx-100) supplemented with protease and phosphatase inhibitor cocktail (Thermo Scientific, USA). Tissues were homogenized for 2x2 min at 50Hz in TissueLyser LT (Qiagen, Germany) with a 0.5 mm metallic ball (Qiagen). Then the tubes were incubated in orbital shaker for 2 h at 4$^{o}$C and centrifuged at 20800 g for 20 min at 4°C. The lysate was transferred to another tube and quantified.

Cells: NRCMs were trypsinized with TripleE, centrifuged at 6000 g for 10 min at 4$^{o}$C to remove the supernatant and the pellet of cells resulted was stored at -80. Then 20 µl of NP40 buffer (Invitrogen) supplemented with protease and phosphatase inhibitor cocktail (Thermo Scientific, USA) was added to each pellet of cells. The pellet was thawed on ice and then frozen again for a total of 5 cycles, with vortexing in between, to completely lyse the cells. Then the tubes were incubated in orbital shaker for 2 h at 4$^{o}$C and centrifuged at 20800 g for 20 min at 4°C. The lysate was transferred to another tube and quantified.

Protein concentrations were estimated by bicinchoninic acid assay (BCA protein assay kit, Pierce). The quantification was performed in 384-well plate (Corning) in a total volume of 50 µl.

Equal amounts of total proteins lysate (20 or 30 µg) were denatured in sample buffer (Invitrogen) supplemented with sample reducing agent (Invitrogen), and electrophoresed on 4-12% gradient Bis-

Tris gels (Invitrogen), with MES SDS running buffer (Invitrogen) and then transferred to 0.2 µm PVDF membrane (BioRad) using BioRad's Trans-Blot Turbo transfer system at 1.3 A and 25 V for 7 min. The membrane was then fixed with 0.4% formalin for 15 min, then washed twice with BPS and blocked with 5% w/v dry milk in 20 mM Tris base, 135 mM NaCl, 0.1% tween 20 (TBST). Membranes were then blotted with primary antibodies indicated in table S4. Blots were later washed and incubated with the respective secondary antibody indicated in table S4. The PageRuler™ Plus protein ladder (Thermo Scientific™) was utilized to estimate the size of the reactive bands. The Western blotting presented in figure 1H was ran on a BioRad system (Laemmli sample buffer supplemented with 10% β- mercaptoethanol, Mini-PROTEAN® TGX Stain-free gels 4-20 %, Precision Plus Protein Dual Color Standards, and Tris/Glycine/SDS for the running buffer, all from BioRad).

Multiple proteins with distinct molecular weights were simultaneously blotted. However, when there was a need to re-incubate a membrane with another antibody, it was stripped with a mild stripping buffer (1.5% glycine, 0.1% SDS and 1% Tween20, pH=2.2) after anti-L2hgdh and D2hgdh blotting, or with a harsh stripping buffer (2% SDS, 62.5mM Tris pH 6.8 and 100mM β-mercaptoethanol) after blotting with OxPhos antibody cocktail. Reactive bands were visualized with SuperSignal West Pico PLUS chemiluminescent substrate (ThermoFisher Scientific) and detected with ChemiDoc™ gel imaging system (BioRad).

Western blotting of isolated mitochondria (**Figure S1F**) was performed using standard protocols. In brief, indicated amounts of mitochondria were solubilized in lysis buffer containing Tris-HCl 60 mM, SDS 2%, glycerol 10%, β-mecaptoethanol 1% and bromphenol blue 0.01%. The cleared homogenate was separated on a 12% SDS-PAGE gel and electrophoretically transferred to a PVDF membrane. Membranes were blocked in TBS containing 5% non-fat dry milk for 120 minutes at room temperature and blotted with the primary antibodies anti-SDHA (tables S4), anti-Vdac, anti-Ndufb8 and anti-Cox4-1 (the latter three antibodies are custom made). Then blotted with secondary antibodies and visualized as it is described above.

The bands were quantified utilizing Image Lab v6.1 (BioRad). The relative band intensity was calculated by dividing each band's intensity on the sum of all bands' intensities of the protein of interest on each blot. The relative band intensity of the protein of interest was then normalized to the relative band intensity of Gapdh on the same blot, and the resulted value was then plotted.

**Table S4: Primary and secondary antibodies utilized in the study.**

| Antibody | Reference number | Provider | dilution | Incubation |
|---|---|---|---|---|
| **OxPhos cocktail** | 45-8099 | Invitrogen | 1:500 | 4ºC, overnight |

| | | | | |
|---|---|---|---|---|
| **IDH2** | MA5-17271 | Invitrogen | 1:1000 | 4ºC, overnight |
| **GAPDH** | MA5-15738 | Invitrogen | 1:2500 | 4ºC, overnight |
| **MLP/CSRP3** | ab155538 | Abcam | 1:2000 | 4ºC, overnight |
| **Succinyllysine** | PTM-401 | PTM Biolabs | 1:500 | 4ºC, overnight |
| **L2HGDH** | 15707-1-AP | Proteintech | 1:200 | 2 h, RT |
| **D2HGDH** | 13895-1-AP | Proteintech | 1:200 | 4ºC, overnight |
| **β-actin** | A5441 | Sigma-Aldrich | 1:5000 | 4ºC, overnight |
| **Cox4-1** | ab16056 | Abcam | 1:1000 | 4ºC, overnight |
| **SDHA** | 5839 | Cell Signalling | 1:1000 | 4ºC, overnight |
| **Secondary anti-rabbit IgG** | NA934 | Sigma-Aldrich | 1:5000 | 1 h, RT |
| **Secondary anti-mouse IgG** | A9044 | Sigma-Aldrich | 1:10 000 | 1 h, RT |

#### 7-2-RNA Extraction

RNA from LV myocardial tissues and NRCMs was extracted by a high throughput method utilizing TRIzol (Invitrogen) following the manufacturer’s instructions with some modifications. In short, NRCMs in each well of the 24-well plate were lysate with 2x75 µl of TRIzol and transferred to 200µl 8-strip tube (Sarstedt). For tissues, 150 µl of TRIzol were added to 2-3 mg of grinded left ventricle cardiac tissues in 200 µl 8-strip tube and pipetted up and down multiple times until the tissues were dissolved. Then after 5 min of incubation at RT, 30 µl of chloroform (Sigma-Aldrich) were added and the resulted two liquid phases were separated by centrifugation. The aqueous phase was transferred to a new 200 µl 8-strip tube containing 1 µl of glycogen (Roche) and the RNA was precipitated overnight by 100 µl of isopropanol at -20 ºC. RNA pellet resulted after centrifugation, was washed twice with 100 µl of 80% ethanol and dissolved in 20-50 µl of RNase-free water. The RNA was quantified with Qubit RNA HS assay (Invitrogen) and utilized for subsequent experiments.

For subsequent applications that required high amounts of RNA with a gDNA digestion step, RNA was instead extracted from 15 mg grinded heart’s tissues. TissueLyser LT (Qiagen, Germany) was used to homogenize the tissue in TRIzol for 2 min at 50Hz with a 0.5 mm metallic ball (Qiagen). After obtaining the RNA pellet, the pellet was washed twice with 80% ethanol and dissolved in 90 µL of RNase free water. Residual gDNA was then digested with the DNase Max® (Qiagen) and the big-RNA fraction (>200nt) was recovered with RNeasy MinElute Cleanup Kit (Qiagen) following the manufacturer’s instructions. The extracted RNA was quantified with Qubit RNA BR assay (Invitrogen) and the quality of RNA was assessed by Fragment Analyzer standard sensitivity RNA kit (Agilent Technologies, USA).

#### 7-3-Reverse Transcription and qPCR

For first strand synthesis of cDNA, SuperScript IV First-Strand Synthesis System (Invitrogen) was used according to the manufacturer's instructions. By utilizing the estimated RNA concentration, the cDNA amount was adjusted to 5 ng/reaction. QPCR was performed using the SYBR® Green Supermix according to the manufacturer's instructions with the following thermal program (enzyme activation at 95°C for 3 min, 40 cycles of [95°C for 10 sec, 60°C for 30 sec, plate read], and followed by recording the melting curve [65 to 95°C, with 0.5°C increment, for 5 sec]). The qPCR was performed in a 384-well plate (BioRad) by utilizing CFX384 Touch™ Real-Time PCR Detection System (BioRad, US). The utilized primers are shown in table (S5). The housekeeping gene Gapdh was used for normalization. All qPCR reactions were run in two technical replicates

**Table S5: qPCR primers utilized in the study**

| Gene | Forward primer | Reverse primer | Species |
|---|---|---|---|
| ***Idh2*** | GGAGAAGCCGGTAGTGGAGAT | GGTCTGGTCACGGTTTGGAA | Mouse & rat |
| ***Hmox1*** | CACGCATATACCCGCTACCT | CCAGAGTGTTCATTCGAGCA | mouse |
| ***Nqo1*** | AGGATGGGAGGTACTCGAATC | AGGCGTCCTTCCTTATATGCTA | mouse |
| ***Osgin1*** | CCTCCGGTATCTGCCTGTC | GGAAAGGTACTCTAGGTCCTGG | mouse |
| ***Gapdh*** | GGGTGTGAACCACGAGAAAT | GTCTTCTGGGTGGCAGTGAT | mouse |
| ***Hmox1*** | GAAGAAGATTGCGCAGAAG | GAAGGCGGTCTTAGCCTCTT | Rat |
| ***Nqo1*** | CTCGCCTCATGCGTTTTTG | CCCCTAATCTGACCTCGTTCAT | Rat |
| ***Osgin1*** | CCTCGGGTATCTGTCTATC | GGAAAGGTACTCTAGGTCCTGA | Rat |
| ***L2hgdh*** | TAGTCATCGTTGGTGGTGGAA | TCCAGTCTGGTGAAGAGCCAAAT | Rat |
| ***D2hgdh*** | GGCTGCCGTTTTCTACCGTGT | GCAGTGCCTAAGAATCTGGGAG | Rat |
| ***Tet1*** | TCCTCAACCCGAGGATGGTA | CTCTTTCCGGGCACACTCAA | Rat |
| ***Tet2*** | TGGTGCTTATGTTCCGTGCT | CAACACCCAGCTCTGTAGGG | Rat |
| ***Tet3*** | CTACATGCCACCACACCACTC | CTGGTTGAGGTTCTTGTGCTG | Rat |
| ***Gapdh*** | GACATGCCGCCTGGAGAAAC | AGCCCAGGATGCCCTTTAGT | Rat |

#### 7-4-Deep RNA sequencing

Prior to RNA-seq, integrity and quantity of the RNA were assessed by Fragment Analyzer Standard Sensitivity RNA kit (Agilent Technologies, USA). All samples had an RNA integrity number >8.7 and were deemed of sufficient quality for mRNA-seq analysis. 1000 ng of total RNA was used as input to each mRNA-seq library. KAPA mRNA HyperPrep kit (Roche, Switzerland) was used for reverse transcription, generation of double stranded cDNA and subsequent library preparation and indexing according to manufacturer's instructions. Quality and quantity of libraries was assessed by Fragment Analyzer standard sensitivity NGS kit (Agilent Technologies, USA). Indexed libraries were pooled in equimolar ratios. The sample pool was quantified with a Qubit Fluorometer (ThermoFisher Scientific, USA) using the dsDNA HS kit (ThermoFisher Scientific, USA), further diluted, and sequenced to >15M paired reads/sample on NextSeq500/550 (Illumina, USA) with PE 75 base pair (bp) read length setting.

Raw RNA sequencing data on myocardium of WT and *Mlp*$^{-/-}$ utilized in figure 6E were obtained from the public repository SRA (SRA accession ID: PRJNA328890). The library preparation and the sequencing are described in the associated metadata on SRA. In brief, high-quality RNA was extracted from whole heart tissues and the libraries were prepared with Illumina TruSeq Stranded Total RNA Library Prep Kit with Ribo-Zero Gold (#RS-122-2301) following the manufacturer's recommendations. Equimolar of the uniquely indexed libraries were pooled and sequenced on the Illumina NextSeq 550 (Paired-end. 75 bp) using sequencing-by-synthesis (SBS) chemistry v4 according to the manufacturer's protocols. Each TruSeq RNA library produced an average yield of 3.6 Gb of sequencing data, with an average of 80% of the reads passing a quality score equal to or greater than Q30. The RNA was from the same heart that was utilized for extracting gDNA for the BS and OxBS libraries, which is described in section (7-6).

**7-5-SmartSeq2 library preparation and sequencing**

Transcriptomic profiling for NRCMs transduced with ShRNA targeting L2hgdh, D2hgdh, Tet1-3 or scramble control was performed by RNA sequencing according to the Smart Seq2 protocol (Picelli et al., 2014), with the help from the Single cell core Facility (SICOF). Each sample was prepared in 4 technical replicates utilizing 25 ng of purified RNA. The mRNA was reverse transcribed into cDNA using oligo (dT) primer and SuperScript II reverse transcriptase (Invitrogen). A template switching oligo was used for the second strand cDNA synthesis. The cDNA was later amplified by PCR for 14 cycles. After purification, the quality of the cDNA was assessed by 2100 Bioanalyzer with a DNA High Sensitivity chip (Agilent Biotechnologies). Then the cDNA was tagmented with Tn5 transposase, and each library was uniquely indexed using the Illumina Nextera XT index kits (Set A–D). The uniquely indexed libraries from half of a 384-well plate were thereafter pooled together and sequenced on one lane of a HiSeq3000 sequencer (Illumina), using dual indexing and single-end 50 base pair (bp) read length setting.

**7-6-Bisulfite and oxidative bisulfite raw data**

Raw data of cardiac whole genome BS and OxBS sequencing on WT and *Mlp*$^{-/-}$ were obtained from the public repository SRA (SRA accession ID: PRJNA328890). The library preparation and the sequencing are described in the associated metadata. In brief, Qiagen DNeasy® Blood and Tissue kit was utilized for extracting genomic DNA from ~15 mg of myocardium of WT and *Mlp*$^{-/-}$ mice (males,12 weeks old, 3 animals in each genotype). TrueMethyl® Whole Genome library preparation kit (Cambridge Epigenetix Alpha Version 1.1, June 2015) was utilized to prepare BS and OxBS libraries according to the manufacturer's instructions. The successful completion of the three distinct procedures; DNA oxidation, bisulfite conversion and post-conversion NGS library construction was assessed through interrogation of the spike-in digestion controls. Equimolar of the resulted 12 uniquely indexed libraries (6 BS and 6 OxBS) were pooled together and then sequenced on 18 lanes of Illumina HiSeq 2500 sequencer (paired-end, 125 bp) using sequencing-by-synthesis (SBS) chemistry v4 according to the manufacturer's protocols. Each library produced an average yield of 91.8 Gb of sequencing data, with an average of 85.4% of the reads passing a quality score equal to or greater than Q30.

#### 8- bioinformatic analysis

##### 8-1- Reads processing of RNA sequencing

Sequenced reads in fastq files were quality controlled and read counts were derived with the bcbio pipeline (1.1.7-b) (Chapman et al., 2020). Reads were aligned to the mouse genome mm10 or to the rat genome Rnor_6.0 using STAR (2.6.1d) (Dobin et al., 2013) with genome annotation from ENSEMBL (version 91). Counts were calculated using featureCounts (1.6.4) (Liao et al., 2014). Differential gene expression analysis was performed with DESeq2 (R package v 1.26.0)

##### 8-2-Epigenetic analysis

For each sample, raw reads of BS or OxBS sequencing from multiple lanes were merged into paired fastq files. The downstream processing analysis was performed according to the TrueMethyl® Data Analysis pipeline (Cambridge Epigenetix (CEGX)). In brief, raw reads were trimmed to remove adapter and low-quality sequence using Cutadapt (v1.11) (Martin, 2011). Trimmed BS and OxBS reads were then mapped to the mouse genome (GRCm38.p4) using Bismark (v0.19.0) (Krueger and Andrews, 2011). The resulting 12 bam files (3 WT, 3 *Mlp* $^{-/-}$, BS and OxBS) were sorted and indexed using SAMtools (v1.8) (Li et al., 2009) and were then passed into the deduplication and methylation extraction tools of Bismark. Base methylation counts were computed from the Bismark output using Bismark methylation extractor tool.

##### 8-2-1 Estimation of 5mc and 5hmC

The average percentage of whole genome 5mC across all biological samples was inferred from the OxBS Bismark alignment report. The whole genome percentage of 5hmC was inferred by the direct subtraction of methylated cytosines' percentages between the BS and OxBS samples from the Bismark alignment report. The percentages of 5mC and 5hmC were found to be 58.7% and 4.4%, respectively. The very low percentage of 5hmC would require a minimum of 100X sequencing coverage to be reliably estimated at a single base pair resolution. Alternatively, an increase in the statistical power can be achieved through pooling of neighboring CpGs with a small compromise in the actual resolution (Booth et al., 2013).

The depth of averaged coverage of publicly available BS and OxBS sequencing data for each biological replicates was estimated to be ~8x. Therefore, a CpGs pooling strategy was developed to boost the minimum coverage to over 100X. Using R (v.3.6), the CytosineReports files were normalized by using methylKit library (v1.12.0) (Akalin et al., 2012) according to the median coverage option. Sliding windows of 30 CpGs with a step size of 2 CpGs were generated for all CpGs in the reference genome (every CpG is covered by 15 windows).

All CpGs with an outlier high coverage (more than the 99.9% percentile) were removed. The "unite" function with (min.per.group=1) was utilized to remove all CpGs that had no initial coverage in any of the sequenced libraries. All remaining CpGs (target-CpGs) in every defined sliding window were tiled together and only windows with a minimum of 100X combined coverage were included in subsequent analysis. Then the pooled window coverage was divided by the number of CpGs and the resulting values was assigned to each CpG inside that window. As each individual CpG would be covered by multiple overlapped windows, the resulting multiple assigned coverages for each CpG were averaged again to obtain the final pooled coverage of each individual remaining CpG. Thereafter, MLML2R (Kiihl et al., 2019; Qu et al., 2013) in (R package v0.3.3) was used to estimate 5mC, 5hmC and non-methylated C at every CpG.

**8-2-2 DMR analysis**

Methylation levels and coverage profiles of every CpG site quantified with the bismark_methylation_extractor tool were further used for Differential Methylation Region (DMR) analysis with the "bsseq" R library (Hansen et al., 2012). The purpose of this analysis is to utilize the natural strong correlation between neighboring CpG sites that typically results in their identical methylation levels. In this way, one can potentially detect long stretches of DNA (beyond individual CpG sites) with differential methylation between two groups of samples. We filtered away CpG sites covered by less than two reads in both WT and *Mlp*$^{-/-}$ samples, as there was not enough statistical evidence for reliable quantification of the methylation levels of those sites. Further, methylation levels of all individual CpG sites were smoothed using the "Bsmooth" algorithm (Hansen et al., 2012) for taking into account the context methylation levels of the neighboring CpG sites. Next, t-statistic was computed in a sliding window across the mouse reference genome with "Bsmooth.tstat", and thresholding on the computed t-statistics was implemented by the "dmrFinder" function in the "bsseq" R library. For the downstream analysis we kept only DMRs with at least three CpG sites and the difference of at least 10% in average methylation levels between WT and *Mlp*$^{-/-}$ for 5mC, and 1% for 5hmC data. This resulted in 26 052 DMRs for 5mC and 138 369 DMRs for 5hmC data. Finally, the identified DMRs were ranked by the sum of the t-statistics of comprising CpG sites weighted by the number of CpG sites in each DMR, i.e. the "areaStat" metric reported by the "bsseq" R library.

**8-2-3 CpG and DMR annotation**

Both individual CpG sites and DMRs were annotated by their overlapping functional elements such as exon, intron, CDS, promoter, enhancer, CTCF, TF binding site, and open chromatin regions. For this purpose, we used mm10 mouse reference genome annotation downloaded from the UCSC resource http://hgdownload.cse.ucsc.edu/goldenpath/mm10/database/, and Ensembl Regulatory Build http://ftp.ensembl.org/pub/release-99/gtf/mus_musculus/. For overlapping CpG and DMR coordinates with the functional regions, we used "bedtools closeset" tool and selected only CpG / DMR – functional region pairs that had the 0-distance implying that they overlapped. Further, for each functional group of elements (exon, intron, CDS, promoter, enhancer, CTCF, TF binding site, and open chromatin regions) we counted the number of times they overlapped the DMRs between WT and *Mlp*$^{-/-}$ identified previously (intersection) and normalized this number by the total amount of DMRs and functional elements (union). This allowed us to construct a type of Jaccard enrichment metric (intersection over union) for investigating what group of functional elements the DMRs were predominantly overlapping with.

**8-2-4 Combining methylation data with RNAseq**

RNA-seq data for *Mlp*$^{-/-}$ and WT samples were aligned with STAR (Dobin et al., 2013) quantified with featureCounts (Liao et al., 2014) and normalized with TMM normalization (Robinson and Oshlack, 2010). Further, for each functional element (exon, intron, CDS, promoter, enhancer, CTCF, TF binding site, and open chromatin regions), DMRs overlapping the functional elements were matched to their closest genes using a custom R script, and methylation levels of the DMRs were correlated against gene expression levels of the corresponding genes using Spearman rank correlation within each WT and *Mlp*$^{-/-}$ sample. Due to a very expensive computational procedure of averaging methylation levels of individual CpG sites falling within DMR regions overlapping functional elements, this was performed on randomly drawn 1000 examples of each group of functional elements.

**8-3-Pathway analysis**

Differentially expressed or methylated genes were analyzed using Ingenuity pathway analysis (IPA: QIAGEN Inc., https://www.qiagenbioinformatics.com/products/ingenuity-pathway-analysis (Kramer et al., 2014)). Differentially expressed genes (DEG) with *p* values below 0.05 were included in the analysis. When the number of DEG were over the permitted limit by IPA, DEG with *p* values below 0.01 were included in the analysis. However, in the IPA comparison analysis, when multiple data sets were compared together, the same threshold of *p* value (i.e. 0.01 or 0.05) was utilized for all data sets included in same comparison analysis. The calculated *p*-value of the IPA core analysis is based on the Right-Tailed Fisher's Exact Test and it reflects the likelihood that an association between a set of significant molecules in the investigated data set and a given pathway is due to random chance. The smaller the *p*-value is, the less likely that an association is random. The activation z-score reflects the likely activation states and its direction for a pathway, compared to a model that assigns random regulation directions. Plotted pathways in all figures were selected from a list of significantly enriched pathways based on their relevance to the context.

#### 9-Statistical Analysis

All data are presented as mean values ± SEM. Unpaired or paired *t*-test was calculated with GraphPad Prism 8 software. If other tests were used, detailed descriptions were included in the figure legends.

#### 10- Reagents

Chemical reagents were of highest grade and obtained from Sigma (Germany) unless indicated otherwise. Table S6 indicates the characteristics of the some of the reagents mentioned in the material and method part.

**Table S6: Characteristics of some of the utilized reagents**

| Material | Notes | Manufacturer | Reference ID |
|---|---|---|---|
| DMEM | DMEM, GlutaMax, 4.5g/L LD-glucose, pyruvate | Gibco | 31966-021 |
| Medium 199 | Contains Eale's salts and L-glutamine | Gibco | 31150-022 |
| Horse serum | | SigmaAldrich | H1138 |
| HEPES | 1M | ThermoFisherScientific | 15630-080 |
| Penicillin/streptomycin | 100X | ThermoFisherScientific | 15140-122 |
| Cytarabine | | SigmaAldrich | C1768 |
| DL-Sulforaphane | | SigmaAldrich | S4441 |
| Gelatin | Gelatin from porcine skin | SigmaAldrich | G1890 |
| RIPA buffer | | Substratenheten Huddinge | MIK2933-1000 |
| Halt™ Protease and Phosphatase Inhibitor Cocktail | 100 X | Thermo Scientific | 78440 |
| Stainless Steel Beads | 5 mm | Qiagen | 69989 |
| NP40 Lysis Buffer | | Invitrogen/Thermo Fisher Scientific | FNN0021 |
| Pierce™ BCA Protein Assay Kit | | Thermo Fisher Scientific | 23227 |
| NuPAGE™ LDS Sample Buffer | 4X | Invitrogen/ Thermo Fisher Scientific | NP0007 |

| | | | |
|---|---|---|---|
| NuPAGE Sample Reducing Agent | 10X | Invitrogen/ Thermo Fisher Scientific | NP0009 |
| NuPAGE™ 4 to 12%, Bis-Tris, 1.0 mm, Mini Protein Gel. | 15-well | Invitrogen/ Thermo Fisher Scientific | NP0323BOX |
| NuPAGE™ MES SDS Running Buffer | (20X) | Invitrogen/ Thermo Fisher Scientific | NP0002 |
| PageRuler™ Plus Prestained Protein Ladder, | 10 to 250 kDa | Thermo Scientific™ | 26619 |
| TBST | pH=7.5 | Substratenheten Huddinge | MIK2334-1000 |
| SuperSignal West Pico PLUS Chemiluminescent Substrate | | Thermo Scientific | 34580 |
| TRIzol™ Reagent | | Invitrogen/ThermoFisher Scintific | 15596026 |
| Chloroform | | Sigma-Aldrich | C2432 |
| Glycogen | | Roche/Sigma-Alrdich | 10901393001 |
| Qubit™ RNA HS Assay Kit | | Invitrogen/ThermoFisher Scintific | Q32855 |
| Nextera XT Index Kit v2 | Sets A-D | Illumina | FC-131-2001,2,3,4 |
| (RS)-2-Hydroxyglutaric acid, disodium salt | (2,3,3-D3; OD, 98%) CP 95% | Cambridge Isotope Laboratories | DLM-9104-PK |
| L2HG standard | | Sigma-Aldrich | 90790 |
| D2HG standard | | Sigma-Aldrich | 16859 |
| TrypLE | | ThermoFisher Scintific | 12604013 |
| isofluran | | Baxter Medical AB | N01AB06 |
| Mini-PROTEAN® TGX Stain-free gels | 4-20 % | BioRad | 4568095 |
| Precision Plus Protein Dual Color Standards | | BioRad | 1610374 |
| Laemmli sample buffer | 4x | BioRad | 1610747 |
| Tris/Glycine/SDS Electrophoresis Buffer | 10x | BioRad | 1610772EDU |

**Table S7: Characteristics of some of the utilized accessories**

| Material | Notes | Manufacturer | Reference ID |
|---|---|---|---|
| Multiply ® -µStrip Pro 8-strip | RNase free | Sarstedt | 72.991.002 |
| SafeSeal micro tube 2ml | | Sarstedt | 72.695.500 |
| Assay plate 384 well | Black with clear flat bottom | Corning | 3683 |
| Hard-Shell® 384-Well PCR Plates, thin wall, skirted, clear/white | | BioRad | HSP3905 |
| Falcon® 6-well Clear Flat Bottom TC-treated Multiwell Cell Culture Plate | | Corning | 353046 |
| Falcon® 24-well Clear Flat Bottom TC-treated Multiwell Cell Culture Plate | | Corning | 353047 |
| Retsch Grinding Ball ss 2 Ø | | Retsch /Germany | 22.455.0010 |
| Corning® 100 mm TC-treated Culture Dish | | Corning | 430167 |
